# Biophysical characterization of high-confidence, small human proteins

**DOI:** 10.1101/2024.04.12.589296

**Authors:** A. M. Whited, Irwin Jungreis, Jeffre Allen, Christina L. Cleveland, Jonathan M. Mudge, Manolis Kellis, John L. Rinn, Loren E. Hough

**Affiliations:** BioFrontiers Institute, University of Colorado, Boulder, CO, USA; Broad Institute of MIT and Harvard, Cambridge, MA, USA; MIT Computer Science and Artificial Intelligence Laboratory, Cambridge, MA, USA; Department of Biochemistry, University of Colorado Boulder, CO, USA; European Molecular Biology Laboratory, European Bioinformatics Institute, Wellcome Genome Campus, Hinxton, Cambridge, UK; Department of Physics, University of Colorado Boulder, CO, USA

**Keywords:** Peptides, Genome, Human, Amino Acids, Proteins, Computational Biology

## Abstract

Significant efforts have been made to characterize the biophysical properties of proteins. Small proteins have received less attention because their annotation has historically been less reliable. However, recent improvements in sequencing, proteomics, and bioinformatics techniques have led to the high-confidence annotation of small open reading frames (smORFs) that encode for functional proteins, producing smORF-encoded proteins (SEPs). SEPs have been found to perform critical functions in several species, including humans. While significant efforts have been made to annotate SEPs, less attention has been given to the biophysical properties of these proteins. We characterized the distributions of predicted and curated biophysical properties, including sequence composition, structure, localization, function, and disease association of a conservative list of previously identified human SEPs. We found significant differences between SEPs and both larger proteins and control sets. Additionally, we provide an example of how our characterization of biophysical properties can contribute to distinguishing protein-coding smORFs from non-coding ones in otherwise ambiguous cases.

**Why it Matters:** The identification of proteins smaller than 100 amino acids has lagged being that of larger proteins in part because their small size makes proteomic and genomic approaches more challenging. However, small proteins can perform significant functions. There are now enough proteins with high confidence of function to enable meaningful comparison with larger proteins, and to determine whether putative small proteins might be identified, based on their biophysical properties. As an example, we found that small proteins often contain transmembrane helices but are relatively devoid of beta sheets, which helped to support the annotation of a putative protein as functional. We also identified a wide range of biological functions and localization for small proteins, including many with disease associations.

## Introduction

Proteins annotation historically focused on proteins longer than approximately 100 amino acids. However, the discovery of small proteins with essential function(1, 2) launched a genetic gold rush to find small open reading frames (smORFs) in the genome of numerous organisms including yeast, bacteria, fruit fly, and other metazoans that are thought to encode functional proteins shorter than 100 amino acids, or smORF-encoded proteins (SEPs). Efforts to identify smORFs and SEPs include mass spectrometry, Ribo-SEQ, and improved conservation scoring algorithms(3–7).

Verified SEPs have been found to serve many functions(6). One of the first SEPs discovered was a peptide of just 11 amino acids translated from a previously annotated lncRNA from a polycistronic gene in *Drosophila* and plays a crucial role in the development of fruit fly legs(1). Additional functional *Drosophila* SEPs with known orthologs in humans were subsequently identified(8). Soon after, evidence of translation of smORFs from other genes previously thought to have only non-coding function were found in metazoans. For example, SEPs were found to be involved in skeletal and cardiac muscle regulation in humans and mice(9, 10). SEPs have been found to produce proteins that serve a diverse set of functions including metabolic homeostasis(11), gene expression and tissue folding(1), cell-to-cell signaling during development(12, 13), phagocytosis(14), embryogenesis(15), DNA repair(16) protein processing(17, 18), and mitochondrial translation(19) and respiration(20).

Genomic characterizations of polypeptide-encoding smORFs have focused on prediction and detection of translation(1, 8–10, 21–25). Such smORFs are found in a myriad of locations within the genome including 5’ and 3’ untranslated regions of genes (UTRs), and within coding sequences (CDSs)(21). A significant challenge in identifying functional SEPs genetically has been the ubiquitous ribosome binding of smORFs(26) potentially with function in the immune system(27–29), and the low conservation of SEPs(25). In addition, there is growing evidence of ubiquitous translation of the genome, including evidence that evolutionarily young sORFs encode for proteins which serve biological functions(30–33). Mass spectrometry approaches, including those with targeted enrichments strategies, have led to the detection of many SEPs in cells (3, 34–38). However, there remains significant uncertainty about the functional significance of putative SEPs. For example, many putative SEPs previously characterized have subsequently been found to have been false positives, instead being pseudogenes, fragments of longer genes, hypothetical translations of non-coding DNA, or regulatory elements within the 5’- UTRs of proteins (See Supplementary Information). As is the case for lncRNAs(39), improvements in analysis tools have and will result in higher fidelity lists of functional SEPs.

We hypothesized that biophysical characterization of high-confidence SEPs can shed light on unique biophysical properties of small proteins and may help distinguish the subset of functional SEPs with evolutionary selection on their biophysical properties. We hypothesized that conservation of biophysical features may be stronger than that of sequence, as is the case for intrinsically disordered proteins(40). As a result, a biophysical lens may help to elucidate proteins with ambiguous annotation. Previous studies, albeit some on lower-confidence datasets (See Supplementary Information), have included such properties as sequence conservation(22), amino acid usage(21, 23), localization(24), and structural properties, e.g. prevalence of transmembrane helices(21, 36, 41, 42) or disorder(23). Structural predictions are available for a wide range of putative and alternative smORFs in the Openprot database(43). In contrast to other studies which have sought to catalogue large permissive sets of candidate SEPs(44, 45), we sought to characterize only high-confidence SEPs. While it is likely that the current annotation of SEPs is missing functional proteins, there are enough high-confidence SEPs to form the basis for biophysical characterization of these proteins.

In this study we have compiled an annotated, manually curated, high confidence, list of 173 human SEPs 16 to 70 amino acids long (HC-SEP). We further reduced this list to remove highly redundant proteins to create a list of 140 “non-redundant SEPs” (NR-SEP). We then developed a biophysical description of these human SEPs using properties that we calculated or predicted from their amino acid sequences or that have been experimentally observed or reported in literature. We characterized expression and diseases association of the HC-SEPs. Finally, we apply these comparisons to a case study to contribute to the annotation of a putative SEP, SMIM42.

## Methods

### Identifying SEPs from GENCODE

GENCODE v37 human transcripts were obtained from the GTF file on the GENCODE web site (https://www.gencodegenes.org). Our initial list of 1330 smORFs consisted of the distinct ATG-initiated coding sequences (CDS) from the 1616 GENCODE transcripts that had biotype protein_coding and whose CDS encoded a protein ≤70 amino acids, excluding transcripts with tags cds_start_NF or cds_end_NF (which indicate incomplete CDS), or for which the length of the CDS was not a multiple of three nucleotides. PhyloCSF scores were calculated using the 58mammals parameter set, the bls option, and the default mle and AsIs options, applied to an alignment of the complete ORF excluding the final stop codon.

Alignments were extracted from the 100 vertebrates whole genome alignments with hg38 reference downloaded from the UCSC Genome Browser(46) and restricting to the 58 placental mammal subset. PhyloCSF-Ψ scores, which correct for the non-independence of neighboring codons, were calculated as described in(4), using coefficients in the file `PsiCoefs.hg38_58.58mammals.mle.txt` from the PhyloCSF installation.

To cut the list of 1330 smORFs down to a size that would be more manageable for manual curation, we excluded 231 with PhyloCSF-Ψ ≤ 0 and most of the 958 that overlapped a longer coding sequence, resulting in a list of 347 candidates for manual curation consisting of:

- 143 that did not overlap a longer coding sequence.
- 153 for which the non-overlapping part of the smORF had PhyloCSF-Ψ > 0.
- 51 for which it was unclear if the unique part of the longer coding sequence was in fact protein-coding (PhyloCSF-Ψ ≤10 or phylogenetic branch length of species in the local alignment was less than half of the whole species tree, so PhyloCSF score might have insufficient statistical power) and either the smORF was entirely contained in a longer coding sequence or the non-overlapping part was too short to confidently call it non-coding (less than 10 amino acids).

We then manually examined the alignments of each of these 347 candidates using CodAlignView (https://data.broadinstitute.org/compbio1/cav.php), checking for conservation of the start and stop codons, lack of frame-shifting insertions and deletions in other species, incorrect alignments, and artifacts that can inflate PhyloCSF score such as overlap with possible enhancer regions. The result is a list of 142 smORFs having clear evolutionary evidence of protein-coding function.

Finally, we added 31 other smORFs for which the PhyloCSF signal was ambiguous or missing but for which there was clear evidence of protein-coding function from mass spectrometry or single-gene studies, resulting in a total of 173 smORFs with high confidence of encoding functional small proteins (high-confidence SEPs, forming the HC-SEP set).

### Refining the list of smORFs and reducing redundancy

Our HC-SEP set included more than one member of each of several distinct protein families. They are: beta-defensins (16), keratin-associated proteins (17), metallothioneins (12), thymosins (6), and guanine nucleotide-binding proteins (6). Since the members of a protein family tend to have similar characteristics, we selected only representative sequences from each of these families to avoid skewing the biophysical characterizations of smORFs in this study.

A multiple sequence alignment of each family using T-COFFEE(47) identified very high levels of similarity (≥95) among metallothioneins, thymosins, and guanine nucleotide-binding proteins. As such, only one representative sequence from each of those families was selected for biophysical characterization in this paper. Gene MT1A encoding for metallothionein-1A, TMSB4Y encoding for thymosin beta-4, and GNG5 encoding for guanine nucleotide-binding protein subunit gamma-5 were chosen as the representative sequences for biophysical characterization of their respective families based on their high conservation scores relative to the other sequences in their families.

The keratin-associated proteins (KAPs) did not exhibit high similarity using T-COFFEE. However, several families and subfamilies of KAPs have been previously identified(48) based on their characteristic sequences. We chose one representative sequence from each predetermined subfamily for a total of eight KAPs for inclusion in the biophysical characterization. For KAP subfamily 22, we used both sequences for the biophysical characterization as they did not align well to each other using EMBOSS Needle for pairwise alignment (identity and similarity below 38% and an alignment score of 73.5). Otherwise, we selected the representative sequence with the highest conservation score using T-COFFEE from each KAP subfamily and a total of 8 KAPs were used.

The largest family of proteins in the HC-SEP set is beta defensins. Based on a multiple sequence alignment using T-COFFEE, there is overall poor similarity of the beta defensins among the HC-SEPs. Furthermore, based on the available literature, there do not exist any clear families or subfamilies of beta defensins. This is unsurprising as the beta defensin genes are amongst the most rapidly evolving mammalian genes(49) and influence the evolution of adjacent genes(50). They have lineage specific duplications which are common alongside rapid sequence divergence. We performed a pair-wise alignment using EMBOSS Needle of all beta defensins in this study to each other and removed one member of any pair with identity of over 95%, reducing the number of beta defensins used for biophysical characterization from 16 to 10.

Present in the set were what appeared to be other protein families based upon their name, e.g. small integral membrane proteins. However, a literature search and multiple sequence alignments of these families revealed that there was no basis for assuming they are distinct protein families unto themselves. Thus, all these smORFs were included in the biophysical characterization.

The result of this pruning was a list of 140 non-redundant SEPs and we refer to this list as the NR-SEP set. The lists of sequences, with those removed and those retained, are available in the Supporting materials. Comparisons in this work are always made between size-matched lists, so that the negative and randomized control sets (described below) are matched to the NR-SEP set.

### Negative and Randomized Controls

We generated three different classes of controls to test different hypotheses about protein function. First, we hypothesized that these proteins would show different behavior than the hypothetical translation of length-matched ORFs that are unlikely to encode functional proteins. To generate this list of “negative control” hypothetical SEPs, we began with every ATG-to-stop smORF of length 16 to 70 codons (not including the stop codon) in the 5’-UTR of a GENCODE protein-coding transcript, excluding ones that overlap an annotated coding sequence or pseudogene in any reading frame on either strand. To assure they were not coding, we required that the PhyloCSF score per codon was less than -18 and that the relative branch length of the local alignment was at least 0.8 (to gain confidence that PhyloCSF had adequate statistical power). We also excluded any whose alignment in any species overlapped the alignment of an annotated (human) CDS or pseudogene, to avoid unannotated pseudogenes. For each of the 140 smORFs in the NR-SEP set, we chose a control smORF of the same length, assuring that no two of these control smORFs overlapped. We then computed the hypothetical translations of these smORFs to create a negative control SEP list.

Second, as the NR-SEPs utilize a slightly different amino acid composition than either UniProt proteins or our negative control set, we sought to determine whether features of these proteins were dictated by this amino acid composition, or whether they were dictated by the ordering of the included amino acids. As such, we generated a “randomized control set of SEPs, by creating a size-matched list of 140, methionine-start sequences whose non-methionine-start amino acids were determined from randomly scrambling the non-start amino acids of the entire pruned NR-SEP set. As a result, the average amino acid composition of the entire randomized control SEP set is identical to that of the NR-SEP set, but the amino acid composition of each protein is variable.

Finally, we sought to determine whether the features of the NR-SEPs were similar to typical, and typically longer, proteins. To compare to known proteins of all lengths, we downloaded the Uniprot UP000005640 reference proteome, and either analyzed the entire proteome (which we called ‘All Uniprot’) or randomly sampled proteins from the proteome to make lists, each of 140 sequences, which we called e.g. Uniprot-1. For every metric tested, the behavior of Uniprot-1 was statistically indistinguishable from the other samplings of the UniProt proteome (e.g. UniProt-2, etc.).

All our sequence sets are available in the Supporting materials.

### **C**haracteristic predictors and analysis

We used PSIPred 4.0(51, 52) or DSSP(53, 54) analysis of AlphaFold(55, 56) structures to predict the secondary structure of the small proteins in this study. Signal peptide prediction was performed using the SignalP-6.0 server(57) for eukaryotes. Transmembrane helices were predicted using the DeepTMHMM Server(58). Cellular localization was predicted using DeepLoc-1.0(59). Disorder was predicted using flDPnn(60). Low complexity regions were identified using SEG(61, 62). Phase separation was predicted using ParSev2 using the calculated values for the entire sequence rather than the window calculations used for longer proteins(63, 64). Default parameters were used for all programs unless otherwise noted.

The grand average of hydropathy (GRAVY) (65) was calculated using MATLAB, with identical results to the online calculator http://www.gravy-calculator.de/index.php. Averaged hydrophobicity using the Miyazawa scale (66) along the sequences aligned from their C-terminus was calculated and smoothed (gaussian smoothing with width of 30) using MATLAB. Cysteine density and charge density for each sequence was calculated using MATLAB or Google Sheets.

GO annotations were compared to the human proteome using PANTHER(67), using the PANTHER Overrepresentation Test (Released 20231017) and the PANTHER GO-Slim comparison sets. The ID list used was the proteins’ UniProt IDs; only 99 of the IDs of the NR- SEP set were mapped to GO annotations.

### Tissue expression and disease association

Using RNA-Seq TPM (transcripts per million) values in GTEx Portal v8(68) (accessed on 2021.05.24), we determined tissue-specific transcript prevalence of the putative smORFs in this study. First, all tissue subtypes were collapsed into one tissue. If a particular smORF transcript was expressed in more than one tissue subtype, we collapsed tissue subtypes into one tissue and took the corresponding mean of the median TPM values for that tissue. Otherwise, the TPM value for a single tissue was used. We used the raw TPM values for expression to make comparisons to three benchmark genes with varying but ubiquitous expression levels. We used the human ribosomal gene RPL18A as a benchmark for overall high and ubiquitous transcription. We used the human protein alpha tubulin, a common control in Western Blots, encoded for by the gene TUBA1A for ubiquitous expression at a mid-range level. Finally, we used the gene for a less common kinesin, kinesin-like protein KIF14, encoded by the gene KIF14, to establish comparison to low level ubiquitous expression.

Many of the proteins not annotated in UniProt did not have TPM values. To assess the pathology of the HC-SEP set, we searched the EMBL-EBI GWAS catalog(69) by genomic location for each transcript, noting pathological SNPs in any part of the transcript. We recorded the p-value and associated trait for each SNP and grouped similar disease states together for a total of 492 p-values in 17 groups. We also searched the OMIM database(70) by gene name for experimentally or clinically validated genetic diseases and recorded the affiliations.

### Statistical comparisons

Nearly all statistical analysis was done using Prism 9 from GraphPad. Data sets were first tested for normality using the D’Agostino-Pearson test and Q-Q plots. While some individual datasets were normally distributed by this test, none of our characterizations had all four datasets being normally distributed. As a result, we used non-parametric tests. We first used the Kruskal-Wallis multiple comparison tests to determine whether at least one sample can be statistically distinguished from the others, and to reduce error inflation. The statistical significance plotted in the figures is the adjusted P values from the Dunn’s multiple comparison test using the Kruskal-Wallis rankings. These pairwise significance values are plotted (ns: p > 0.05, *: p ≤ 0.05, **: p ≤ 0.01, ***: p ≤ 0.001, ****: p ≤ 0.0001).

Each control set was chosen to test a specific hypothesis, and so we additionally used Mann-Whitney test (aka Wilcoxon rank sum test, performed in Prism 9 for all but Figure 2 and in MATLAB 2021, for Figure 2) to test the significance of pairwise comparisons between sets with continuous variables and the Fisher’s Exact test (performed in MATLAB) for comparisons between binary data (e.g. contains a signal peptide or not)(71). All uncertainties listed in the text are standard error of the mean. All p values are listed in Table S1.

## Results

### Identifying High Confidence SEPs

To obtain the most accurate possible characterization of biophysical properties of SEPs, we used evolutionary signatures and manual curation to generate a high confidence list of smORFs that encode functional proteins (Fig. 1A).

**Figure 1.**
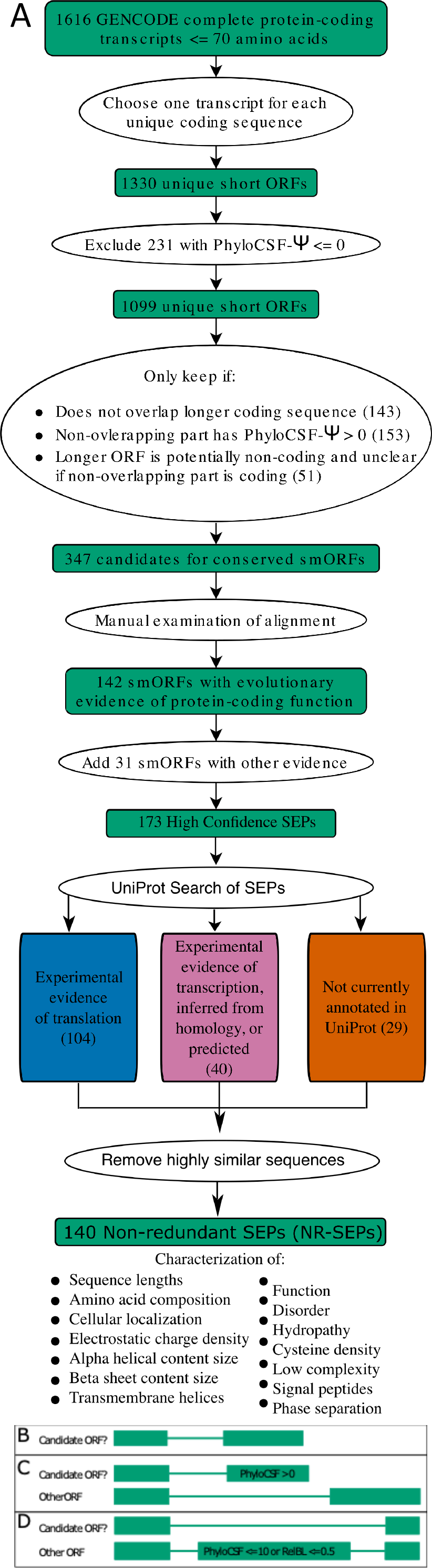
Schematic of the procedure to generate and analyze the HC-SEP and NR-SEP lists. (A) A flow chart shows the steps in generating the SEP datasets characterizing their biophysical properties. (B-D) The positive PhyloCSF score of an ORF could be due to an overlapping protein-coding sequence rather than the ORF itself. To avoid this possibility, we restricted to those ORFs with positive PhyloCSF score satisfying one of three conditions to generate a list of 347 candidate ORFs which were then manually evaluated. The three conditions were: (B) ORFs that do not overlap another annotated coding sequence. (C) ORFs that partly overlap another annotated coding sequence, but which contain a non-overlapping portion with positive score. (D) ORFs that are entirely or almost entirely contained in another annotated coding sequence that lacks a strong protein-coding evolutionary signature so the longer ORF might not actually be protein-coding.

**Figure 2.**
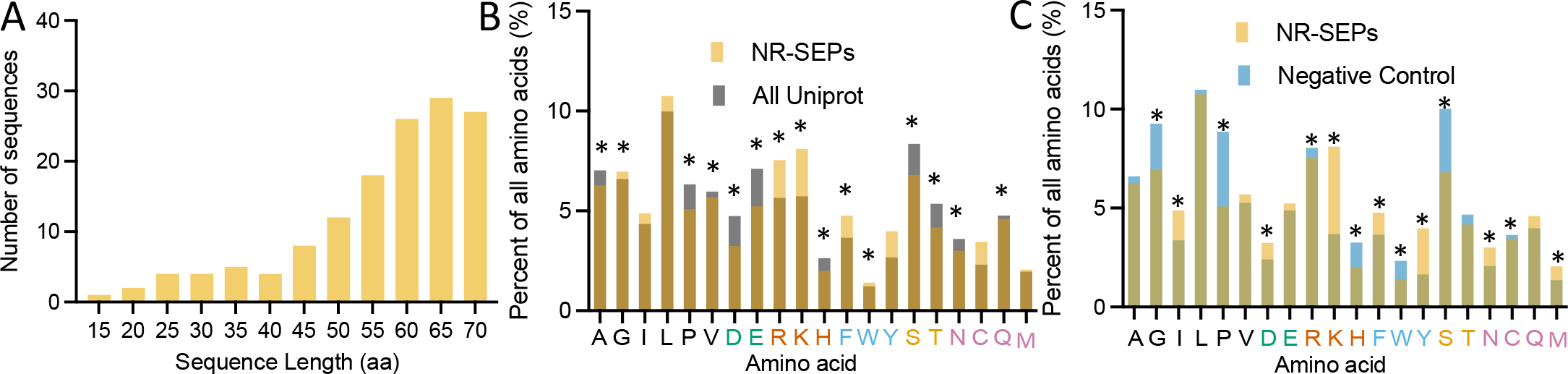
Analysis of the lengths and amino acid composition of SEPs. (A) A size distribution of the 140 NR-SEPs shows a skew towards the high end of the range X-labels are bin centers. (B) A comparison of the amino acid frequencies of the NR-SEPs (gold) to the amino acid frequencies of the proteins in the UniProt database (grey). Amino acids labels are colored by approximate property: aliphatic (black), acidic (green), basic (red), aromatic (blue), hydroxylic (gold), amidic or sulfur containing (purple), and start methionines were excluded from the analysis. The NR- SEPs are enriched in arginine, lysine, and cysteine and depleted in negatively charged amino acids glutamate and aspartate. (C) A comparison of the amino acid frequencies of the NR-SEPs (gold) and the negative control (blue). The negative controls are enriched in serine, proline, and glycine while the NR-SEPs are enriched in lysine and tryptophan. Significance (*) indicates p<0.05 in the Mann-Whitney test comparing the distribution amino acid density within proteins in each set (Table S1).

We began with the 1330 complete ATG-initiated smORFs no more than 70 amino acids long that are annotated as protein-coding in GENCODE(72) version 37. However, some of these were annotated as protein-coding in the early 2000’s when fewer resources were available and would no longer be considered protein-coding based on current evidentiary standards.

Furthermore, once a gene has been shown to be protein-coding, GENCODE is permissive in assigning protein-coding status to other transcripts in that gene even without strong evidence that the particular transcript encodes a functional protein. Consequently, we imposed many additional filters and manually examined the multi-species alignment of every candidate, to obtain a subset of smORFs that we can be confident encode functional SEPs. The importance of this additional filtering and manual curation is highlighted by a previous study of biophysical properties of SEPs that used a collection of ORFs <= 100 aa that were predicted to be protein-coding based on evolutionary conservation(23). Although the prediction method produced a fruitful ranked list of novel candidates, the vast majority of the novel predictions have since been determined not to be protein-coding ORFs, limiting their usefulness for distinguishing properties of SEPs. In particular, after manual curation by GENCODE of the 831 predicted novel protein-coding ORFs, only 37 were considered likely to be true protein-coding ORFs. Hundreds were pseudogenes, others were partial matches to longer protein-coding ORFs, and the majority were false discoveries due to the large number of ORFs tested (Supporting material).

We first calculated PhyloCSF-Ψ(4) scores for each of the 1330 GENCODE smORFs and excluded smORFs having score less than zero. PhyloCSF compares codon substitution frequencies in alignments of closely related genomes to coding and non-coding evolutionary models trained on whole genome data to distinguish genomic regions that have evolved under protein-coding constraint, and PhyloCSF-Ψ is a log likelihood ratio indicating how much more or less likely a protein-coding genomic region of a particular length is to get a particular PhyloCSF score than a non-coding region. However, while a positive PhyloCSF score provides evidence that a *genomic* region is protein-coding, that does not prove that a particular transcript is protein-coding because the genomic region can be shared by several transcripts. In particular, a positive score could result from an ORF part of which is protein-coding and part is not, due to overlap with the coding sequence of a different splice variant. To address this, we first excluded smORFs that overlap a longer coding sequence unless the unique part of the original smORF has positive score, or the portion of the longer coding sequence that does not overlap the original smORF lacks a strong evolutionary signature of protein-coding function (Fig. 1B-D). We then manually examined the alignment of each of the remaining 347 ORFs using CodAlignView to find 142 for which the evolutionary signature showed clear evidence of protein coding function that was not due to overlap with a different protein coding transcript.

Finally, we added 31 other smORFs for which the PhyloCSF signal was ambiguous or missing but for which there was clear evidence of protein coding function from mass spectrometry or single-gene studies. Most of these were paralogs of known protein-coding genes that were poorly aligned in the multispecies alignment, including 12 keratin associated proteins and 7 beta-defensins, though there were also rapidly evolving genes that did not have a PhyloCSF signal. The resulting high-confidence SEP set (HC-SEP set) included 173 ORFs encoding small proteins ranging from 16 to 70 amino acids in length.

We performed a UniProt search of the full HC-SEP set(73). Using the level of annotation and review in UniProt, each HC-SEPs fell into one of three categories: genes with experimental evidence of translation; genes that have experimental evidence of transcription, homology to a known protein-coding gene, or identified as potentially protein-coding in bioinformatic studies; and genes that were not annotated in UniProt at the time of our search (April 2022, Fig. 1A).

This is a similar approach to that of Ma et al(34). We found that 104 of the proteins have direct experimental evidence at the protein level, and 40 more are annotated in UniProt, typically based on translation inferred from homology or prediction. Finally, 29 proteins were unannotated in UniProt at the time of our curation.

As the HC-SEP list included several sequences from each of several families with very high sequence similarity, we further reduced the list by picking a representative sequence from each of these groups. We reasoned that otherwise the characterized biophysical properties would be skewed by this redundancy. This reduced the number of keratin-associated proteins, beta defensins, metallothioneins, thymosins, and guanine nucleotide-binding proteins. The result was 140 proteins which we call the “non-redundant SEP” set (NR-SEP). The pruned list contains 79 proteins that have direct experimental evidence at the protein level, 32 more that are annotated in UniProt, and 29 that were unannotated in UniProt at the time of our curation.

We also generated a list of 140 non-coding, ATG-initiated “negative control” smORFs in 5’-UTRs of protein-coding genes that have strongly negative PhyloCSF score, do not overlap a protein-coding ORF or pseudogene in any frame, and that match the lengths of the 140 NR-SEP smORFs. Any instances of translation from the 5’-UTR, as occasionally seen(31), may dilute our negative control set, however, any difference we find significant between the positive and negative controls with this dilution will become more significant in comparing true positive and negatives. We then generated a negative control set of hypothetical SEPs by the hypothetical translation of the negative control smORFs. This negative control set allowed us to identify distinguishing features of SEPs as compared to the hypothetical translation of otherwise similar genetic locations.

We created an additional control set of 140 sequences (“randomized controls”), size matched to the NR-SEP set, by scrambling the total amino acid content of the NR-SEP set and maintaining a methionine start for each protein. This randomized control set allowed us to distinguish those characteristics of small proteins that are consequences of their amino acid bias as opposed to independent sequence features.

Finally, we compared either the full set of proteins in the UniProt UP000005640 reference database (one protein per gene), or randomly selected six lists of 140 non-redundant sequences from UP000005640 for comparison (UniProt-1 to UniProt-6) when comparing to the full UniProt set would have been computationally prohibitive. Figures generally contain either the full UP000005640 set or UniProt-1, as the results were consistent between the different UniProt-1 to UniProt-6 subsets (Fig. S1).

### Characteristics of SEP sequence properties Sequence length distribution

The mean size of the NR-SEPs in this study is 57 amino acids (Fig. 2A). The smallest is just 16 amino acids. Overall, there is a bias towards longer SEPs, with 62% of the NR-SEPs between 60-70 amino acids in length.

### Amino acid composition

We compared the amino acid composition of the 140 NR-SEPs to the amino acid composition of proteins included in the *Homo sapiens* UniProt reference proteome (UP000005640) (Fig. 2B) and the amino acid composition arising from the hypothetical translation of the negative control smORFs (Fig. 2C). There was no comparison to the randomized control as its overall amino acid composition is identical the amino acid composition of the NR-SEPs. As we required a methionine start codon which biases the prevalence of methionine, we included only non-start methionines in the analysis.

The amino acid prevalence of the NR-SEPs is different from those in the human UniProt database (p values from the Wilcoxon rank sum test are given in Table S1). The most striking differences between the NR-SEPs and UniProt proteins was in the preference for positively charged amino acids (arginine and lysine, though not histidine) and depletion of negatively charged amino acids (both aspartate and glutamate) in the NR-SEPs. The NR-SEPs also appear to utilize fewer serine and threonine residues than the UniProt database. This could suggest that small proteins may undergo less phosphorylation than their canonical protein counterparts. In contrast, the NR-SEPs are enriched in the aromatic residue phenylalanine relative to UniProt proteins. The NR-SEPs are also depleted in proline in comparison to the UniProt database and a depletion of proline residues affects the protein structures that can form(74).

There is a slight increase in cysteine utilization by the NR-SEPs relative to UniProt proteins, with 3.5±0.4% of all residues being cysteines in the NR-SEPs set, but only 2.54±0.02% in UniProt (p=0.08). This result is largely a result of three protein families, the beta-defensins, keratin-associated proteins, and metallothioneins. Of proteins within the HC-SEP set, the average cysteine density for metallothionein is 32%, for keratin-associated proteins is 11%, and for beta-defensins is 9%. Excluding these family members from the NR-SEPs gives a mean of 2.23±0.3%. The amino acid frequencies of the NR-SEPs appear different from the amino acid frequencies obtained by hypothetical translation of the negative controls. As with the comparison to the UniProt database, the NR-SEPs show the strongest enrichment in lysine and tyrosine residues. The NR-SEPs are notably depleted in glycine, proline and serine when compared to the negative controls.

### Cysteine density

Given their importance in secondary structure formation, we characterized the cysteine density of SEPs to determine if a higher cysteine prevalence occurred uniformly across the set, or if only a subset of proteins contained a high density of cysteines. The cysteine density of the NR- SEPs, negative controls, randomized controls, and the UniProt database was calculated as percentage of the overall sequence by counting the number of cysteine residues in the sequence and dividing by the sequence length (Fig. 3A).

**Figure 3.**
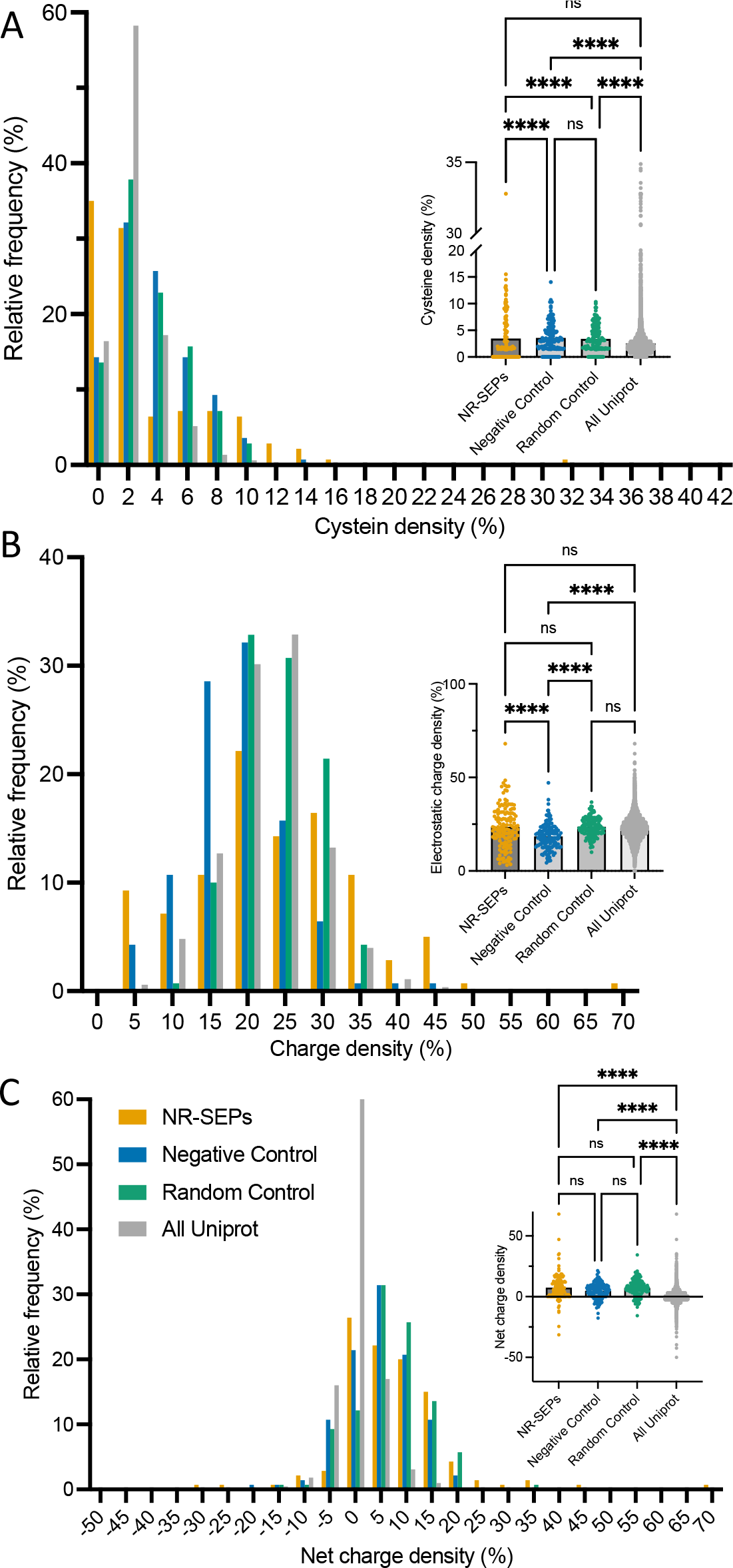
Biophysical characteristics of SEPs calculated from their amino acid sequences. (A) NR-SEPs have the highest fraction of proteins lacking cysteines, and a high overall cysteine density. NR-SEPs have the broadest distribution of (B) charge densities and (C) net charge density. In addition, the NR-SEPs have an average net positive charge, in contrast with UniProt where the mean net charge is near zero. (x-labels are bin centers, * indicates statistical difference with a p value of < 0.05, Table S1)

The NR-SEPs have a larger fraction of proteins with no cysteines and lower median cysteine density than any of the other datasets, even though they have a higher mean cysteine content than the UniProt database. Moreover, the standard deviation of cysteine density is highest for the NR-SEP set. In other words, cysteines are more unequally distributed among the NR-SEPs than the other datasets. These observations of uneven cysteine density among the NR-SEPs are consistent with the very high cysteine density among the beta-defensins, keratin-associated proteins, and metallothioneins. Pairwise differences between NR-SEPs and the negative or randomized controls are all statistically significant (Table S1). The cysteine density distribution of the NR-SEPs is quite different from that of the UniProt set, with the NR-SEPs having both more ORFs with 0 density and more with very high density (6% or greater) but fewer with intermediate density (2-4%), so the Rank Sum test does not find either one to significantly dominate the other.

### Electrostatic charge density

The over- and under- representation of positively or negatively charged amino acids respectively led us to characterize the electrostatic charge density of SEPs. We calculated both the charge density, by counting the total number of positively and negatively charged residues in the sequence and dividing by the sequence length, and the net charge density, as the length normalized difference between the number of positive and negatively charged residues.

The distribution of charge densities of the NR-SEPs appears broader than the other datasets, with a standard deviation almost twice that of the other distributions (roughly 11 vs 5 or 6). The distribution of charge densities of the NR-SEPs neither significantly dominates nor is dominated by the distribution for the randomized controls (as expected) or the distribution for the UniProt dataset (Fig. 3B, Table S1). Electrostatic interactions are important modulators of protein interactions and structures(75) and are responsible for protein-protein interactions, protein function, and hydration and they may be responsible for some of the unique features of small proteins.

The net charge density of NR-SEPs was much more like that of the two control sets than of proteins in UniProt. UniProt proteins had a median net charge of zero (mean of 0.3±0.03), and a relatively narrow distribution (standard deviation of 4.2), whereas the NR-SEPs mostly have positive net charge, with a median net charge of 6 (mean of 7.4±0.9) and a standard deviation of 10.9. This net charge is significantly different even from that found in most organisms(76).

### Hydropathy

We characterized the hydropathy of SEPs using the grand average of hydropathy (GRAVY) value. This value is determined by summing the Kyte-Doolittle hydropathy values of all amino acids in the sequence and dividing by the sequence length(65). Positive values indicate hydrophobicity and negative values indicate hydrophilicity. None of the data sets significantly dominates any of the others except that the random controls have slightly higher values than UniProt proteins (Fig. S2, Table S1). The NR-SEPs have a broader distribution, with more proteins showing extreme hydrophobicity and hydrophilicity than any of the other datasets.

Indeed, the standard deviation is almost twice as large for the NR-SEPs as for UniProt proteins (0.83 vs 0.43). For purely statistical reasons, one would expect the average value of any property to be more variable among shorter proteins than among longer ones, but the fact that GRAVY scores are more variable among NR-SEPs than among the random controls, which have the same lengths and even the same amino acid frequencies, implies that the tendency of NR- SEPs to have either very high or very low hydropathy is biologically meaningful, rather than a statistical artifact.

Recently, charge near the C-terminus of proteins was identified as a distinguishing feature of conserved proteins as compared to the ubiquitous translation of potential ORFs(77).

We plotted a smoothed Miyazawa hydrophobicity(66) scores for the different sets and found a general decrease in hydrophobicity moving towards the C-termini for both the UniProt and NR- SEP sets but not for either of the control sets, as expected.

### Characteristics of SEPs predicted from their amino acid sequences Secondary structure prediction

We used PSIPred to predict the secondary structure of the NR-SEPs, negative controls, and randomized controls(Figs. 4, S1) (52). Because PSIPred could readily be used on putative proteins constituting our control datasets, we used it as the primary method of comparison. In addition, we used DSSP to extract secondary structure from AlphaFold structures of the NR-SEP and Uniprot-1 sets to additionally validate any findings from the PSIPred analysis (53–56). Our main findings were recapitulated using AlphaFold predicted structures (Fig. S2, S3). Eight of the sequences in the NR-SEPs and each of the size-matched control sets were too small (28 amino acids or less) for secondary structure prediction by PSIPred and those were removed from the analyzed pool of SEPs. We additionally computed secondary structure predictions for 6 sets of 140 proteins randomly sampled from UniProt (Uniprot-1 to Uniprot-6) with no significant difference seen among those 6 sets for any metric we considered (Fig. S1).

**Figure 4.**
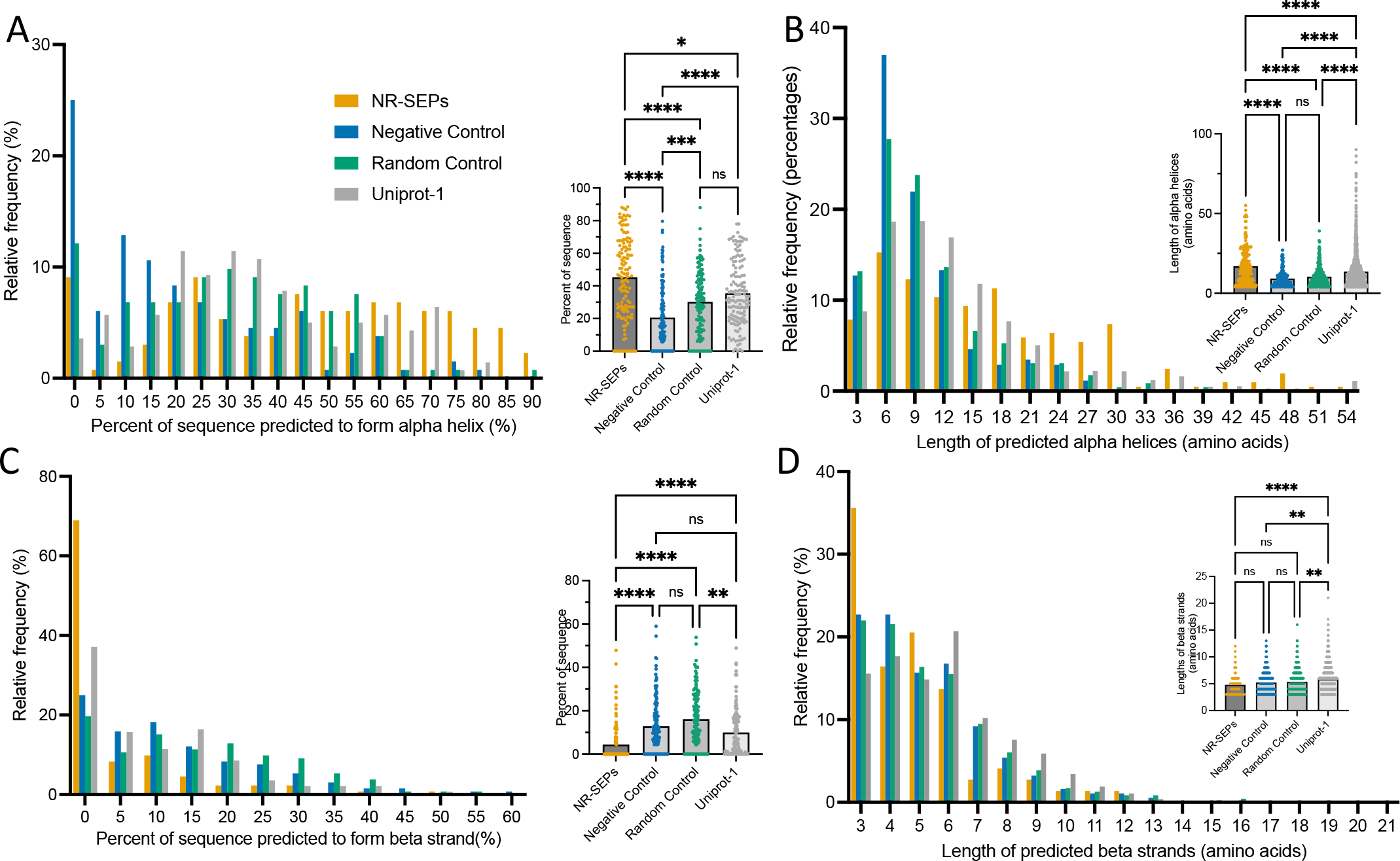
Predicted structural biophysical characteristics of SEPs. (A) The NR-SEPs contain more predicted alpha helical content than the controls. Relative to UniProt proteins, more NR- SEPs lack any alpha helical regions (10 vs 1), but those that do have helical regions have a higher alpha helical content. (B) The helices are predicted to be larger in the NR-SEPs than the controls. All helices larger than 54 were collapsed to the final bin in the histogram for ease of viewing. (C) The NR-SEPs contain lower fraction of predicted beta strand content than the control and of proteins in Uniprot, although the predicted strands are of similar size to the controls. (x-labels are bin centers, * indicates statistical difference with a p value of < 0.05, Table S1)

We required four consecutive amino acids for alpha helices and three amino acids for beta strands for them to be considered in our analysis. The percentage of the sequence occupied by a predicted secondary structure was calculated by counting the number of amino acids predicted to form a particular secondary structure and dividing it by the length of the sequence.

Secondary structure prediction is a key feature of this study as broad structural characterization remains a large gap in the smORF literature. There are substantial differences in the PSIPred predicted secondary structures of the NR-SEPs and the controls (Fig. 4A, Fig. S1). Despite containing more proteins with no predicted alpha helical content than the UniProt set, the NR-SEPs contain far more predicted alpha helical content than all three controls (significance in Table S1), with a mean fraction of 45±2% of amino acids predicted to be in an alpha helix of at least 4 amino acids. For comparison, the mean fraction of the UniProt-1 set was 35±2% (very similar to the other Uniprot sets, Fig. S1). Moreover, the predicted alpha-helical length was longest in the NR-SEPs, with a mean length of 17±1 amino acids, as compared to 13.7±0.3 amino acids for the UniProt-1 set (Fig. 4B). The average length of proteins in the UniProt-1 set is 580±30 amino acids, giving the UniProt-1 proteins much longer sequences with which to make longer alpha helices. In contrast, the NR-SEPs tend to have a single, long helix, often a transmembrane helix (see below).

Our analysis of the predicted beta strand content of SEPs shows the opposite trend of the alpha helices (Fig. 4C, Table S1). Almost 70% of the NR-SEPs do not contain any PSIPred predicted beta strands whereas only about 20% of the other set do not contain any and the mean fraction of beta strands is significantly smaller for the NR-SEPs than for all other sets, with only 4.5±0.8% of the residues being predicted to form a beta strand, significantly less than e.g. the roughly 10% of residues in the UniProt sets, or 16±1% of residues in the randomized control. Beta strands are thus significantly excluded from these small proteins when compared to either real proteins, or randomized controls containing the same amino acid composition. The distribution of sizes of the few predicted beta strands is statistically similar to those of the negative and randomized controls, but significantly smaller than that of the UniProt-1 (Fig. 4D).

We repeated the comparison of the NR-SEPs and UniProt-1 proteins using DSSP to extract the secondary structural features from AlphaFold-predicted structures. Five proteins in the NR-SEP set and two in the UniProt-1 set did not have AlphaFold-predicted structures and were skipped. As with PSIPred, AlphaFold predicts more and longer alpha helicies in the NR- SEP set than in the UniProt-1 set, with a lower prevalence of beta strands (Figs. S2, S3).

### Disorder prediction

The dearth of beta strands in our analysis led us to consider whether disordered regions were more prevalent in SEPs that in other proteins. The percent of the sequence found to be disordered was predicted using flDPnn(60). The NR-SEPs contain more predicted disorder than either the randomized control or UniProt protein sets, though a similar distribution to the negative controls (Fig. 5A, Table S1). Nearly 8% of NR-SEPs proteins are predicted to be at least 95% disordered, as compared to roughly 1% of the Uniprot-1 sets of proteins, which more typically contain multiple protein domains, not all of which are disordered. The mean fraction of disorder is highest in the negative controls (39±3%), followed by the NR-SEPs (30±2%), UniProt (18±2%) and random controls (18±1%).

**Figure 5.**
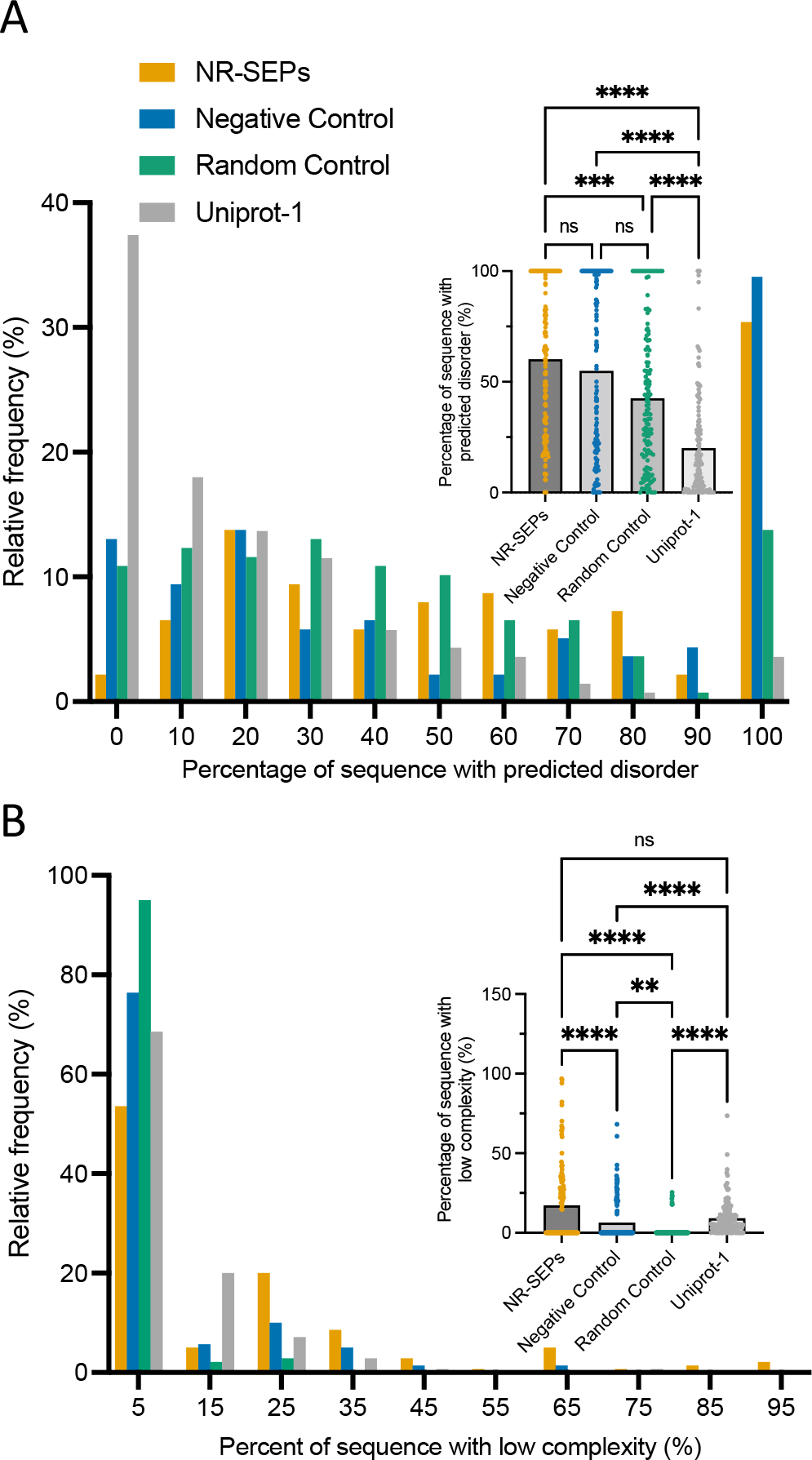
Predicted disordered and low complexity of SEPs as a percent of overall sequence. (A) Predicted disorder was calculated as a percentage of the overall sequence. The NR-SEPs contain more predicted disorder than proteins in Uniport and random sequences, but not than the negative controls. (B) The NR-SEPs contain similar low complexity than proteins in Uniprot, and more than the controls sets. Additional plots of this data are in Supplemental Figure S5, demonstrating that the percent of sequence obscures somewhat the depletion of low percentages of the NR-SEPs, random, and negative control sets because of the minimum size of low- complexity region. (x-labels are bin centers, * indicates statistical difference with a p value of < 0.05, Table S1)

### Low complexity

Low complexity regions (amino acid sequences that contain repeats of single amino acids or short amino acid motifs) have been indicated in the formation of young, novel genes(78) with unique functional domains that enhance protein-protein interactions(79), and solubility and folding(80). Moreover, we had noticed that several of our sequences had large stretches containing primarily a single amino acid. These observations led us to identify the prevalence of low complexity sequences using SEG through the web server PlaToLoCo(61, 62).

A majority of the sequences in the NR-SEP, negative control, and randomized control sets do not contain any identified regions of low complexity (Fig. 5B, S5, Table S1). As such, the median percent of sequence identified as low complexity for all three data sets is zero. As the algorithm used has a minimum size cutoff for low complexity regions (the lowest in this study was 8 amino acids), comparisons to the UniProt set are a bit skewed at low percentage relative to other sets because of their longer lengths. The UniProt-1 set contains a higher fraction of sequences with some low complexity, though a lower fraction of residues in low complexity regions than the NR-SEPs. The NR-SEPs have a larger range than either control and contain more residues in regions of low complexity. In total, 17±2% of residues in NR-SEPs are annotated by SEQ as belonging to regions of low complexity, significantly higher than that for the human genome(61), or the UniProt-1 set (9±1%). By comparison, the average fraction of low complexity in SEPs in the randomized controls is 1.1±0.4% and 6.5±1.1% in the negative controls.

### Transmembrane helices

As we had noticed several long helices in the NR-SEPs, and many of the recently discovered SEPs found experimentally to be translated encode for transmembrane helices (TMHs)(8, 10, 14, 20), we sought to determine the prevalence of transmembrane helices in our sets. Transmembrane helices were predicted using the DeepTMHMM(58). We recorded whether a SEP was predicted to form a TMH or not and then determined the total percentage of SEPs within a data set that form TMHs. A substantial fraction, 30%, of the NR-SEPs are predicted to form TMHs whereas none of the negative or random controls are predicted to do so (Fig. 6A).

**Figure 6.**
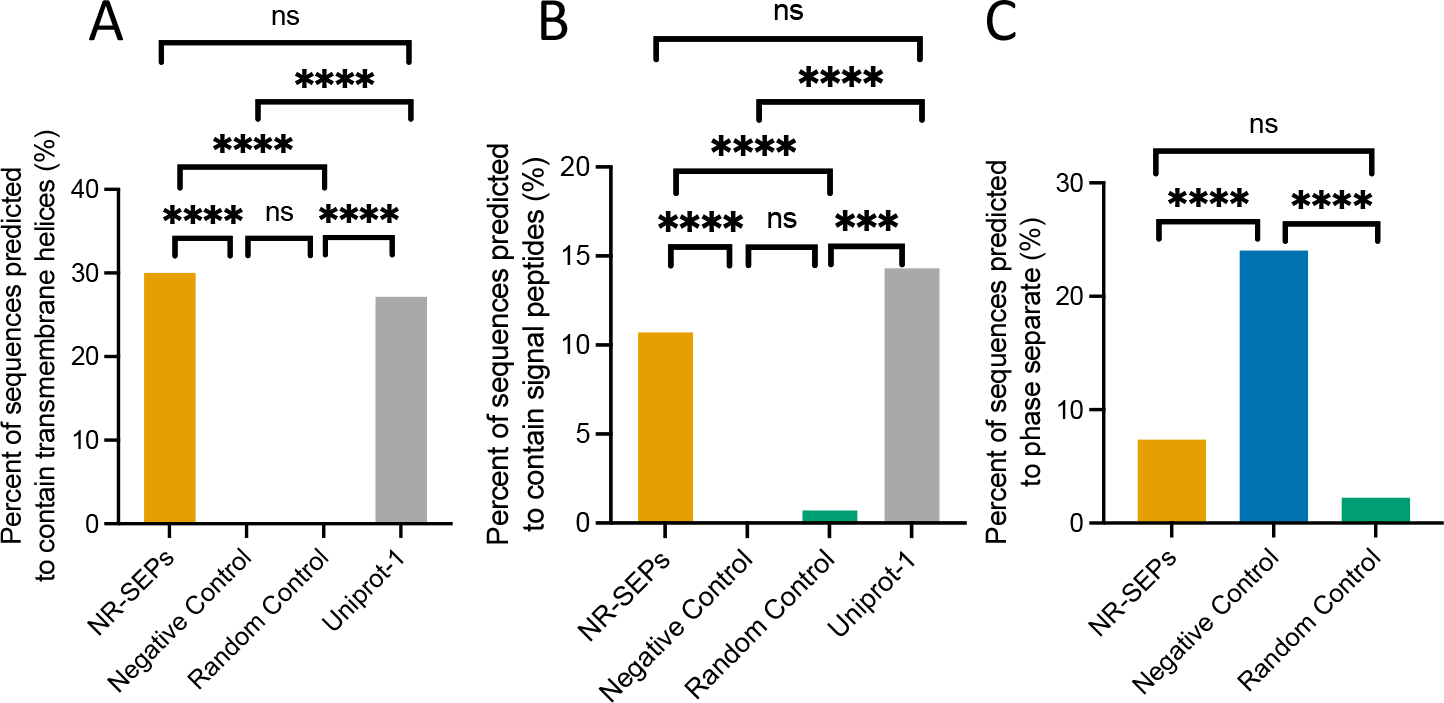
Predicted fraction of sequences containing (A) transmembrane helices, (B) signal peptides, and (C) phase separation propensity. In the first two, NR-SEPs are more similar to proteins in Uniprot than the control sets. All statistical comparison are using the Fisher test (Table S1).

The fraction of NR-SEPs predicted to form TMHs is slightly higher than the fraction of proteins in the UniProt-1 dataset, but the difference is not statistically significant (p=0.7, Table S1). Thus, SEPs are similar to other human proteins in the prevalence of transmembrane helices, which shows a strong bias towards this type of domain given their smaller size relative to UniProt proteins. This is consistent with observations that transmembrane helices are more prevalent in canonical ORFs than alternative ORFs(81).

### Signal Peptides

Adjacent to several of the observed transmembrane helices in the NR-SEPs were signal peptides directing the peptide to its membrane localization. We sought to determine if signal peptides were more generally present. Signal peptides were predicted using SignalP-6.0 for eukaryotic organisms which predicts the presence of three of the most common types of signal peptides(82). We recorded whether a SEP in any data set was predicted to contain a signal peptide and then determined as a percentage how many of the SEPs in a particular data set were predicted to contain a signal peptide. There are far more predicted signal peptides, 11%, in the NR-SEPs, than in either the random or negative control SEPs, which have less than 1% (Fig. 6B). In contrast about 14% of proteins in the UniProt-1 to -6 datasets contain signal peptides by this predictor, with 16.7% in the full reference UniProt set as calculated by the SignalP authors(57).

### Phase separation

Phase separation is of growing importance in our understanding of biology(83). Given that many protein domains that drive phase separation are relatively short(84), SEPs could play a role in biological processes that drive phase separation. Phase separation was predicted using ParSe v2(63). The propensity of a protein to phase separate appears to depend more strongly on the presence of a domain thought to drive phase separation than on the fraction of residues predicted to drive phase separation(64). As a result, only comparisons between similarly sized proteins are meaningful, and so we did not include in our comparison proteins from UniProt, which are longer. The NR-SEPs were significantly less likely to be predicted to phase separate by ParSe than the negative controls (p<0.0001, Fig. 6, Table S1), though similar to the random controls, as expected as ParSe calculates propensity to phase separate based on amino acid composition.

### Comparison between well-characterized SEPs and those unannotated in UniProt

Although there is strong evidence that each of the NR-SEPs is translated to a functional peptide, only 56% of them have direct experimental evidence of translation reported in UniProt. As a sanity check on our results, we compared the distributions of the biophysical properties of those NR-SEPs having experimental evidence of translation reported in UniProt (79 / 140) to those that are not currently annotated in UniProt (29 / 140). The distributions appear visually similar, and in no case did we find one distribution to statistically significantly dominate the other (Fig. S6).

### Experimentally determined characteristics of SEPs

To complement the quantitative comparisons to longer proteins or to control sets, we additionally manually curated experimentally determined features of SEPs, including cellular localization, protein function, and disease associated annotation. Our aim was to determine the range of properties and functions that could be encoded in such small proteins.

### Cellular localization

To determine protein localization, we manually curated experimentally determined or predicted localization given in the UniProt database (Fig. 7A). If there was more than one cellular location associated with a SEPs, all were counted in the analysis. Some NR-SEPs have no localization information reported in UniProt and those were not considered in the annotated analysis. We found that the NR-SEPs in this study are primarily localized within the cell, with 15% secreted. Secreted small proteins have also been seen in experimental studies that used Ribo-Seq and mass spectrometry(85, 86). The most populated cellular compartments containing NR-SEPs in this study are the mitochondrion (16%), nucleus (15%), cytosol (14%), and cytoskeleton (14%). SEPs are also found in lower numbers in the endoplasmic reticulum (ER) (6%), plasma membrane (5%), and the Golgi (0.7%).

**Figure 7.**
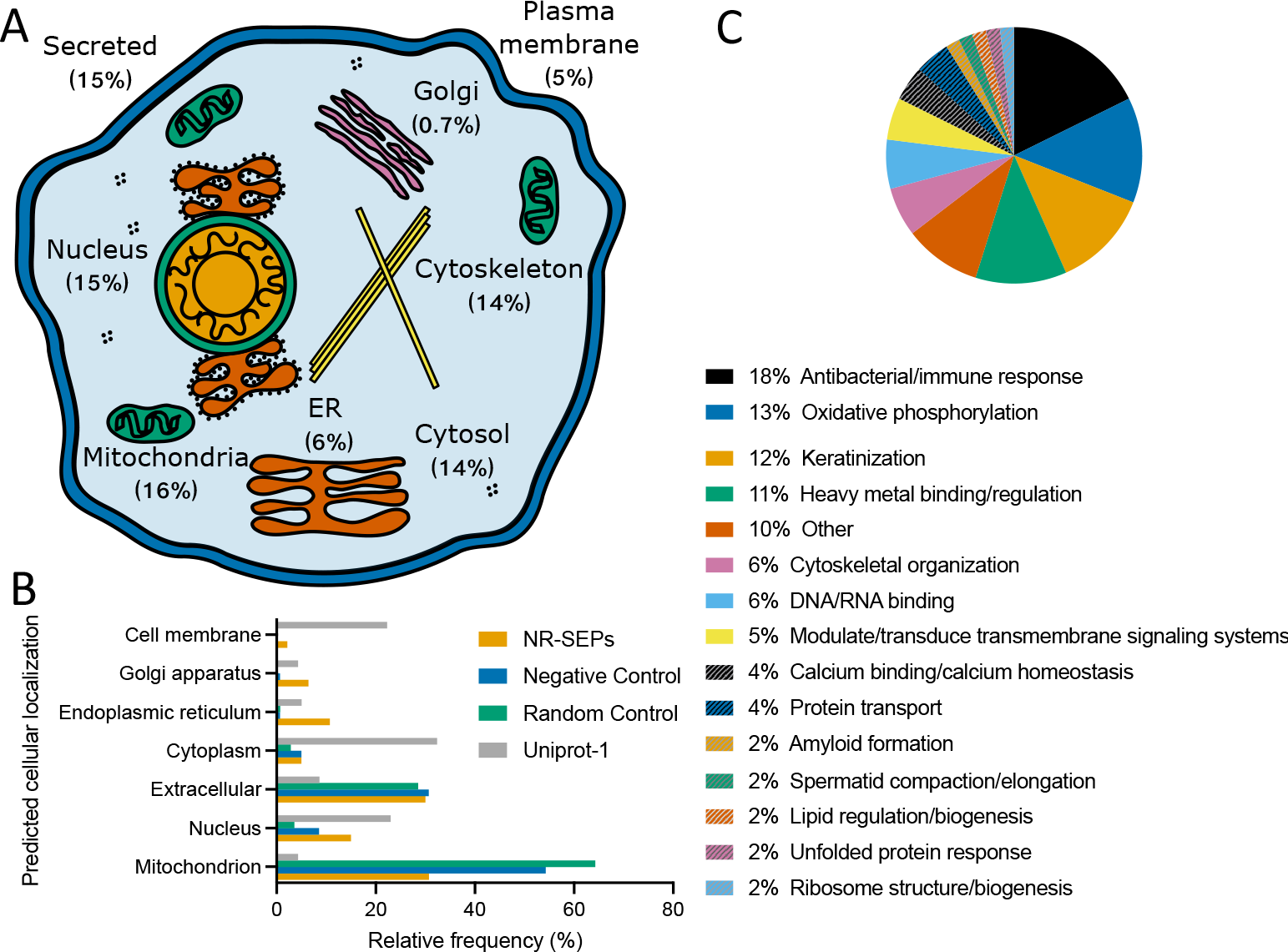
Known and predicted cellular localization of SEPs and their associated functions. (A) The NR-SEPs with annotated cellular localizations in UniProt and literature were analyzed showing a broad and an even distribution through cellular organelles. (B) Predicted, rather than experimentally determined, localization shows a wider range of localization for both NR-SEPs and Uniprot sequences than for control sequences. (C) The NR-SEPs with annotated function in UniProt and the literature were categorized. The most populous functional category is for SEPs involved in antibacterial/immune functions followed by the SEPs involved in the mitochondrion- driven process of oxidative phosphorylation.

Using DeepLoc 1.0(59) to predict cellular localization, we were able to compare the cellular localization of all of the protein sets (Fig. 7B). The NR-SEPs are predicted to be distributed throughout the cell just as seen for the subset of NR-SEPs with annotated cellular localizations. The two most commonly predicted locations of NR-SEPs are the mitochondrion and the extracellular space. However, there are fewer SEPs predicted to reside in the nucleus and cytoplasm than are expected based on the NR-SEPs localizations that have been determined.

This could be a consequence of the limitations on using cellular localization predictors for SEPs. In particular, previous studies have shown that SEPs may contain novel motifs for mediating localization (87), making it difficult for predictors, which have not yet incorporated data from SEP localization experiments (3), to project where in the cell the SEP may reside. There could also be physical consequences to the size of these proteins and their cellular localization. For example, because of their small size, they could potentially bypass physical barriers to larger proteins. In the case of MRI-2, the amino acid sequence does not contain a canonical nuclear localization sequence, yet it is found in the nucleus in addition to the cytoplasm, a fact that has been assigned to MRI-2 being a protein of sufficiently small size that it can freely diffuse through the nuclear pore complex(16).

In contrast, most of the negative and randomized controls are predicted to reside in the mitochondrion or to be secreted. There is very little predicted distribution outside of those organelles, particularly for the randomized control.

To compare the localization of the NR-SEPs to the human proteome, we used PANTHER(67) to determine the statistical overrepresentation of annotated cellular components. 99 of the NR-SEPs were associated with GO annotations. The primary category with significant over-representation relative to the human proteome was the mitochondria (Table S2).

### Function

We combined curated UniProt annotation with manual, literature-based curation to functionally characterize the SEPs in this study that have a known function. For this characterization, we did not remove any of the redundant SEPs as we wished to survey the entire level of representation of each function, and instead used the full HC-SEP set. The largest functional group found in this study is involved in an antibacterial/immune response (Fig. 7C). This is consistent with the high number of beta-defensins in the HC-SEP set and the known prevalence of small antimicrobial proteins found in directed functional studies(88). The HC-SEP set also includes several other antibacterial/immune response proteins that are not beta-defensins. The second largest function group is oxidative phosphorylation which aligns well with our observation that many HC-SEPs are either found experimentally or predicted to be localized to the mitochondrion. The third largest functional group is involved in keratinization and all the members of this functional group are keratin-associated proteins.

Other functional groups include but are not limited to DNA and RNA binding, calcium homeostasis, cytoskeletal organization, and spermatid compaction and elongation. Of the 10% of putative proteins that don’t readily cluster (‘other’), functions include ciliogenesis, cardiovascular development and regulation, and visual perception. It is also worth noting that 35% of the NR-SEPs have a currently unknown function which when determined may lead to further diversity of function beyond what has been found in this study.

To compare functions to the human proteome, we used PANTHER(67) to compare the annotated biological processes of the human proteome as compared to the 100 NR-SEPs that have GO annotations. We found significant overrepresentation of protein involved in regulation of metabolism and significant underrepresentation of proteins involved in transcription and transcriptional regulation (Table S3).

### Tissue expression

Using the available tissue-specific RNA-Seq data in GTex Portal v8(68), we characterized the transcript expression of the full list of 173 HC-SEP ORFs. Shown in Fig. 8 is the TPM value (average among tissue subtypes with measured expression). We used RPL18A, TUBA1A, and KIF14 for comparisons of high, medium and low-level expression. As recommended by the GTex curators, we did not attempt a quantitative analysis because GTex data is not normalized in such a way as to enable such comparison(89).

**Figure 8.**
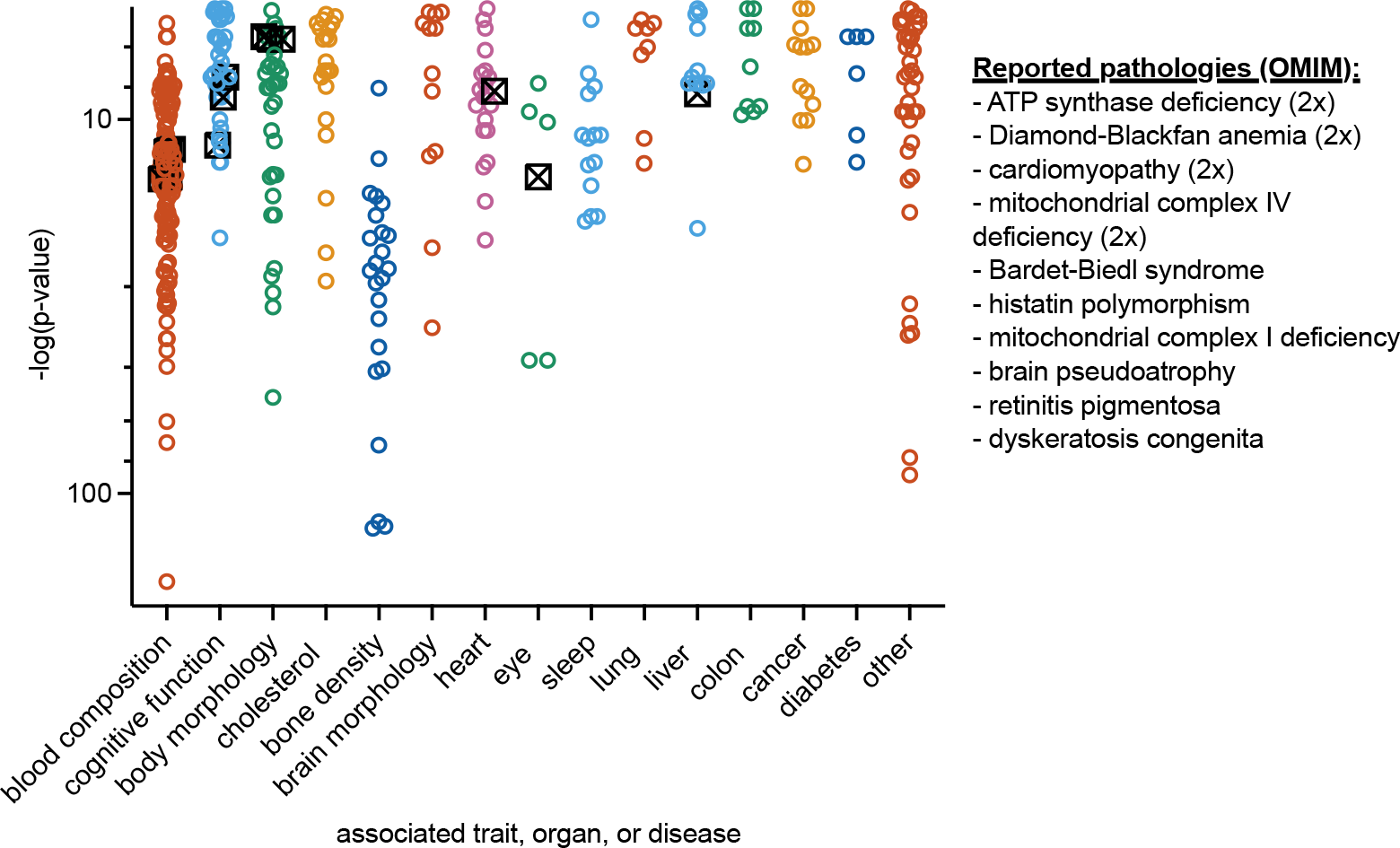
RNA-seq expression data of the HC-SEPs smORFs. Tissue subtypes were collapsed and the mean of the tissue subtype medians of the TPM value for that tissue was used. Several SEPs have high levels of ubiquitous expression and the two tissues expressing the largest number of HC-SEPs are the brain and testis. Some SEPs were not reported to be expressed by GTex (though this is likely due to version and methodological differences between GTex and GENCODE). Included are expression references for RPL18A (high ubiquitous expression), TUBA1A (mid-range ubiquitous expression), and KIF14 (low ubiquitous expression).

Several HC-SEP ORFs are transcribed heavily. The transcript with the highest overall expression level is MT2A which encodes metallothionein 2, a zinc-binding protein, which is known to be expressed in almost all tissues (Fig. 8). The two tissues that express the highest number of HC-SEP ORFs in this study are the brain and testis, similar to ORFs of longer proteins(90, 91).

### smORFs are important for disease

We searched for any pathological single-nucleotide polymorphisms (SNPs) for each of the 173 transcripts from the HC-SEP set in the EMBL-EBI GWAS catalog(69) to determine the impact of genetic variation of small proteins on disease phenotypes. We additionally identified the subset of those SNPs that reside in the coding region. We then categorized each SNPs’ p-value by its associated trait, organ, or disease and plotted the negative log of the p-value (Fig. 9). We did not include any p-values larger than 1e-6 as these are not included in the EMBL-EBI catalog nor are they deemed statistically significant. We found a total of 417 SNPs, 15 of which are in the protein coding region (Fig. 9, Supplemental Materials).

**Figure 9.**
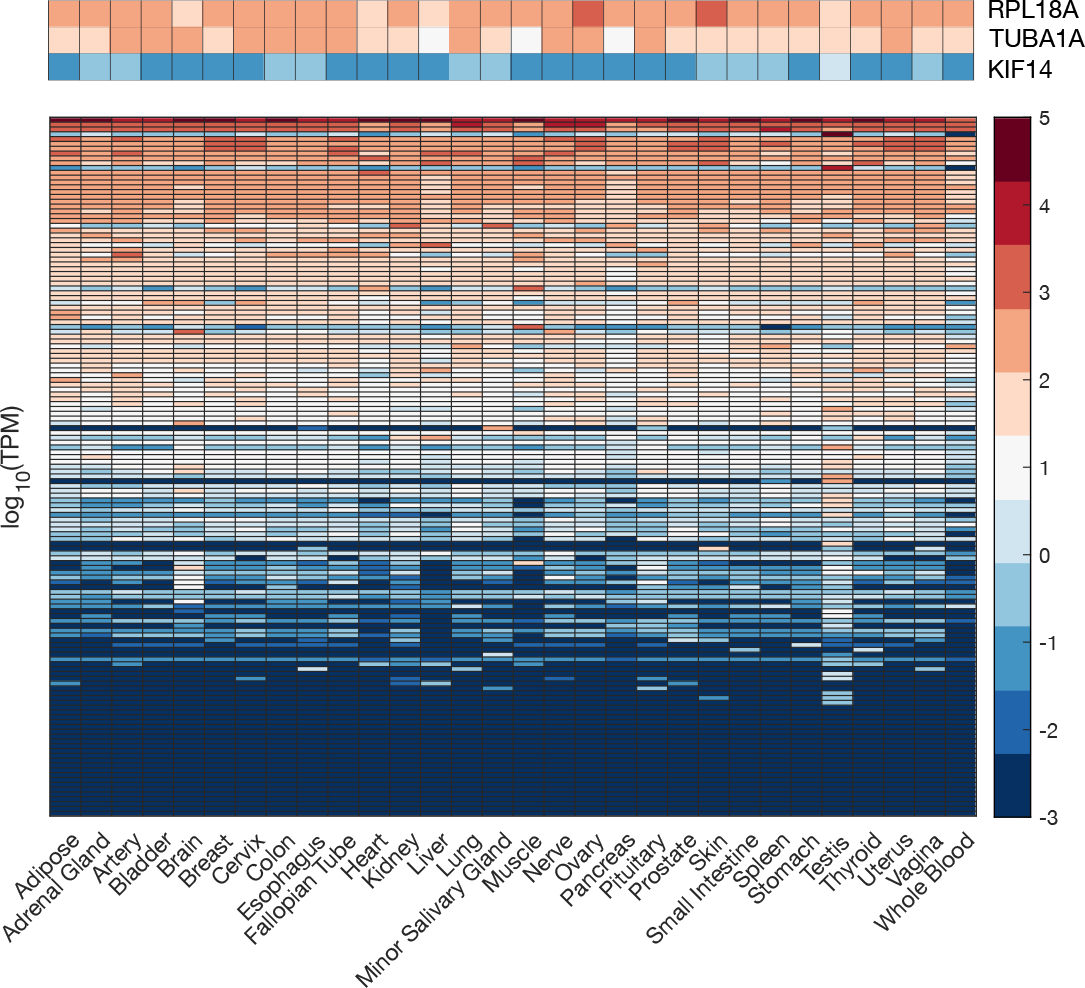
GWAS and OMIM data on smORFs. There have been 417 reported GWAS SNPs within the HC-SEP ORFs (colored open circles). Of those, 15 are SNPs in the coding regions that encode the HC-SEPs (black crossed square). Additionally, 13 of the HC-SEP ORFs are associated with disease in the OMIM database.

Additionally, we searched the OMIM database(70) for experimentally or clinically verified diseases caused by genetic variations in our list of HC-SEPs and reported the clinically or experimentally validated disease and the number of associations for each (Fig. 9, Supplemental Materials). There are a total of 14 reported pathologies for ten different diseases. Interestingly, 5 of 14 (36%) are deficiencies in electron transport chain complexes and are types of mitochondrial complex deficiencies. We conclude that smORFs are important for a wide range of diseases.

### An example: SMIM42

The potential utility of our characterization of biophysical properties of small proteins for helping to distinguish protein-coding smORFs from non-coding DNA is demonstrated by the example of SMIM42, which encodes a 70 amino acid SEP.

Although SMIM42 has been annotated as protein-coding for many years, GENCODE annotators have struggled with the question of whether to remove it because of limited and ambiguous evidence of protein-coding function. There is ample evidence of transcription, with RNAseq datasets supporting testis specific expression (not shown), but there is no mass spectrometry evidence of translation for this protein in Peptide Atlas (Human Protein Atlas proteinatlas.org, Uhlen 2015, (92)). The open reading frame is conserved in many placental mammals but is also lost in several lineages. Among the lineages having intact ORF, the PhyloCSF score is near 0 -- higher than expected for a non-coding region but lower than expected for a protein-coding region. Because of these uncertainties, especially it’s low PhyloCSF score, we had excluded it from our list of high confidence SEPs.

To help resolve this ambiguity, we investigated the biophysical properties of the predicted protein, and found that DeepTMHMM predicts a transmembrane helix twenty-one amino acids in length. Since 30% of the NR-SEPs but none of our negative or random controls are predicted to contain TMHs, this provides support for the conclusion that SMIM42 is protein-coding. GENCODE hopes to incorporate DeepTMHMM predictions and other biophysical properties as additional sources of evidence when making protein-coding determinations in the future.

## Discussion

We characterized high confidence SEPs and compared them to both negative control sets and larger proteins. Our results show that this class of proteins shows robust differences from longer proteins by several measures, and are clearly distinct from control sets, making them a unique protein class. The SEPs differ from longer proteins by several metrics, including their higher net charge density, longer alpha helices, dearth of beta strand, and increased disorder.

The differences in protein secondary structure features are presumably due to the additional restraints of folding such small proteins and could additionally be influenced by limitations on the roles of such small proteins. For example, they are unlikely to be enzymes on their own. It may be that amino acid conformations typically stabilized by beta-sheet structures are more typically stabilized by disulfide bonds, given that some protein families in the SEPs have very high cysteine densities. The high charge density in SEPs may mean that these proteins are more sensitive to cellular pH and ionic strength, which can be modulated in cellular stress and disease(93). The NR-SEPs are similar to longer evolutionarily conserved proteins in that they tend to be depleted in hydrophobic amino acids at their C-termini(77).

Our list of NR-SEPs differ from negative and randomized control SEPs in many ways. It is possible that some of our negative control SEPs are, in fact, functional proteins. If so, these would serve to dilute our comparisons, implying that the statistical differences we observe here would be even stronger in a comparison of the NR-SEPs with a list of true negative controls.

The most notable differences between the NR-SEPs and the controls is in the prevalence of transmembrane helices, signal peptides, and higher cysteine density; these differences might prove useful in the future to help distinguish functional SEP-encoding smORFs from candidates that are not under evolutionary selection(77). The prevalence of transmembrane helices and canonical signal peptides appear to be very strong indicators that a putative ORF is translated into a functional SEP. As an example, we considered the case of SMIM42, which contains a predicted transmembrane helix. We had not included SMIM42 in our list of HC-SEPs, in large part because of its ambiguous PhylosCSF score. However, the prediction of a clear transmembrane helix gives some confidence to the annotation of SMIM42 as a functional protein.

Several experimental and computational limitations may have introduced bias into our list of HC-SEPs relative to the complete set of functional SEPs in the human genome.

Characterizing smORFs that encode functional proteins among the millions of putative smORFs has remained challenging(94) . Experimental limitations in determining which SEPs are functional include losses in mass-spectrometry approaches, and ambiguity of whether RNA or protein products are biologically relevant(30, 33, 77, 95). As a result, our list of well-annotated SEPs may contain biases from these experimental limitations, including potential under- representation of transmembrane proteins. In addition, we imposed sequence requirements which may exclude functional SEPs, including the requirement for a canonical ATG start codon (10, 81, 96–102), and conservation at the codon level, which may bias our set against evolutionarily young proteins(30).

Particularly confounding is the ubiquitous ribosome binding and translation of the transcriptome, which appears to enable rapid response to environmental stimuli(30, 33, 77). Our analysis largely misses the resulting evolutionarily transient proteins, and is primarily of evolutionary conserved SEPs. In addition, it may be that SEPs with certain properties or belonging to certain protein families might have been more amenable to discovery and annotation, leading to a bias in our characterization. As a result, our characterization may not encompass the full range of potential characteristics of all functional SEPs.

There are additional caveats to our biophysical characterization because the predictors used were typically developed for larger proteins. The smallest of the SEPs are too small for many secondary structure predictors including PSIPRED, the secondary structure predictor used in this study. Additionally, many structure predictors rely on the depth of multiple sequence alignments which may be generally lacking for small proteins and is problematic in small protein families such as the beta defensins and keratin-associated proteins in this study that have poor sequence alignment. Similarly, predictors for cellular localization, signal peptides, and cellular functions which may also not be optimized for SEPs which may contain unique, as-of-yet-unknown signaling or recognition sequences(87).

In conclusion, we characterized the biophysical characteristics of SEPs that we identified in a conservative search of the human genome that have a high likelihood of being functional proteins. Our results contribute to an understanding of how small proteins differ from larger ones. These differences may also prove useful for predicting translation and for understanding other aspects of polypeptide-encoding smORFs including their origin and evolution.

## Abbreviations

Small open reading frames (smORFs), smORF-encoded polypeptides (SEPs), long noncoding RNAs (lncRNAs), open reading frame (ORF), ribonucleic acid sequencing (RNA-Seq), ribosome sequencing (Ribo-Seq), untranslated regions of genes (UTRs), coding sequence (CDS), grand average of hydropathy (GRAVY), transmembrane helices (TMHs), endoplasmic reticulum (ER), deoxyribonucleic acid (DNA), ribonucleic acid (RNA), Genome-wide association studies (GWAS), Online Mendelian Inheritance in Man (OMIM). Datasets included in this study are abbreviated HC-SEPs for the list of 173 high confidence SEPs, and NR-SEPs for the subset of 140 proteins curated to reduce redundancy.

## Supporting Material description

The supplement consists of an excel file containing all sequence lists,

## Availability of data and materials

All datasets used and/or analyzed during the current study are included as supplementary information. All analysis are available from the corresponding author on reasonable request.

## Authors’ Contributions

AW, LH, and JR conceived of the project, AW, JA, and CC (supervised by LH), LH, JM, and IJ (in collaboration with MK) designed and performed research for the paper, AW, LH, IJ, and JA prepared figures, AW and LH wrote the manuscript, with input and contributions from IJ, JM, and JR. All authors read and approved the final manuscript.

## Supporting information

Supplemental discussion and figures

List of sequences

