## Supplemental discussion and figures for "Biophysical characterization of high-confidence, small human proteins"

#### Supplementary Section 1

In this section we explain why the novel smORFs predictions by Mackowiak et al (Mackowiak et al. 2015) included so many that were not in fact protein-coding ORFs, and make some recommendations to improve the specificity of future searches for novel smORFs, based on our experience over the last 18 years using PhyloCSF and other comparative genomics methods for distinguishing protein-coding genomic regions.

Mackowiak et al. used the PhyloCSF Omega Test (similar to dN/dS) (Lin et al. 2011; Lin 2012) and other conservation metrics to predict protein-coding function for 1230 non-overlapping human ORFs <= 100 aa, 399 of them previously annotated and 831 novel, as well as many short ORFs in other species, and analyzed properties of these predicted small ORFs. However, extensive manual evaluation of the 831 predicted novel human ORFs by GENCODE found that only a small fraction were in fact protein-coding. Evolutionary methods such as PhyloCSF are extremely powerful for detecting conserved protein-coding regions, but there are pitfalls that need to be avoided. Here we review the methods used and explain why so few of the novel predictions proved correct. We focus on the human predictions, though most of the methodical issues apply to the other species as well.

Mackowiak et al. used a support vector machine (SVM) to score every ORF at least 27 nt in every annotated transcript at protein-coding and lncRNA loci (including transcripts of type NMD, retained intron, etc.) in GENCODE version 19 as well as several other lincRNA catalogs, excluding ORFs that overlap a conserved annotated coding sequence in the same frame or that are contained in one of their other smORFs. The SVM used four features, namely the PhyloCSF Omega score, which was found to be the dominant feature; a measure of frame conservation; and measures of how well the nucleotide-level conservation profiles near the start and end match those of annotated short ORFs. They also found translation evidence for 45 of the novel smORF predictions using published ribosome profiling data sets, and 34 using published mass spectrometry data sets.

We compared the 831 novel human smORF predictions from Mackowiak et al. Table S1 (those satisfying `nonoverlapping=TRUE` for which `smORF_type` was not `annotated` or `ncRNA`) to the annotations in the latest GENCODE release, version 38, which contains many protein-coding regions and pseudogenes that were not present in version 19, including several dozen added as a result of the Mackowiak et al. analysis (Mudge et al. 2019).

We found that the GENCODE annotators considered only 37 of these 831 smORF candidates to be complete protein-coding ORFs. Another 49 partially match a protein-coding ORF, 7 of which are <= 100 aa. This exposes one of the challenges in using PhyloCSF to discover novel protein-coding ORFs, namely that PhyloCSF scores an alignment of genomic DNA, and a region of genomic DNA can be contained in many different transcripts -- if any of these transcripts is protein-coding then the DNA will evolve under protein-coding selection and PhyloCSF cannot directly distinguish which transcripts are responsible. As the authors themselves noted, “incomplete transcriptome assembly could lead to fragmented lincRNA identifications that obscure the presence of longer ORFs”.

GENCODE v19 was released in 2013, and many additional transcripts have been discovered since that time, so the set of transcripts used in the Mackowiak et al. analysis was incomplete. In many cases, the Mackowiak et al. ORF did identify a protein-coding genomic locus, but it was not an smORF, or was not the exact smORF they specified.

We found that 286 of the Mackowiak et al. smORF predictions overlap regions that are now annotated as pseudogenes rather than protein-coding genes. There are another 85 for which part of the smORF and some other annotated coding region are aligned to the same region in one or more of the species in the vertebrate alignment, indicating possible homology and suggesting that the smORF could be part of an unannotated pseudogene. Although Mackowiak et al. excluded ORFs that were annotated as pseudogenes, those annotations are incomplete, and were more so in 2013. This exposes another of the challenges in using PhyloCSF, namely distinguishing protein-coding regions from pseudogenes. PhyloCSF uses an alignment to determine if a genomic region has evolved more like its coding model than its non-coding model. If many, but not all, of the genomes in the alignment have evolved under protein-coding constraint, PhyloCSF will often consider this to be a protein-coding alignment. Regions that are protein-coding in most of the species tree but were pseudogenized in the lineage of the reference species (i.e., unitary pseudogenes) will often get positive PhyloCSF scores. Duplicated and processed pseudogenes can also get positive PhyloCSF scores due to non-syntenic alignment to the protein-coding parent. Mackowiak et al. found that inclusion of measures of nucleotide-level conservation profiles near the start and end of the ORF improved the ability of the SVM to distinguish pseudogenes as compared to PhyloCSF alone; however, they did not include pseudogenes in the negative training set, so their SVM was not specifically trained to distinguish pseudogenes, and they do not report a false positive rate for pseudogenes. ORFs in pseudogenes are particularly likely to be false positives in searches for *short* ORFs, because premature stop codons and frame-shifting indels will often cut a longer ORF into shorter ones.

Mackowiak et al. used a statistical test to determine that most of their smORFs in 3'-UTRs are not parts of longer coding regions that are translated by stop codon readthrough, but did not check individual ones. The authors noted, “A very small number of cases [of 3'UTR smORFs] could be explained by read-through...” In fact, three of the smORFs are in the extensions of the known stop codon readthrough genes *MPZ*, *OPRK1*, and, *OPRL1*.

Evolutionary methods such as PhyloCSF and dN/dS will often find a false signal of protein-coding evolution on the opposite strand from a true coding region in the reading frame that shares the third codon position with the true coding region because a preponderance of substitutions in this position will correlate with a protein-coding signal on both strands. Eight of the Mackowiak smORFs are in this “antisense” frame from true coding regions on the opposite strand. In order to avoid such false discoveries, we recommend excluding from consideration ORFs that have lower PhyloCSF score than the same region on the opposite strand in this antisense frame.

The remainder of the 831 novel smORF candidates are likely to be false discoveries that obtained a high PhyloCSF score due to chance. Mackowiak et al. used cross validation to estimate the false positive rate of their algorithm to be between 0.1% and 0.5%, but noted that, "even with low estimated false positive rates we expect a significant number of false discoveries, since the size of the non-coding transcriptome likely far exceeds the amount of true coding sequence." Based on the description in their methods section, we estimated that they applied their algorithm to approximately 750,000 ORFs, which would be expected to result in between 750 and 3750 false discoveries, so it is not surprising that the bulk of their 831 candidates were false discoveries.

Because there are several distinct ways that an ORF can fail to be a complete protein-coding short ORF (pseudogenes, antisense, partial match to a longer transcript, or just non-coding DNA), we recommend against relying on a single test and instead have distinct tests that distinguish true coding smORFs from each class of alternatives. One must also not rely too heavily on an assumption that existing annotations are complete, since it is precisely the missing splice variants, pseudogenes, etc., that are likely to confound searches for novel coding regions.

The needle-in-a-haystack nature of whole-genome search for novel coding regions dictates that most of the results of even a highly-specific method are likely to be false discoveries. It is therefore essential to perform additional checking for each individual novel candidate. For conserved coding regions, we recommend manually checking the alignment, which can be done using the color-coding of protein-coding-relevant features in CodAlignView (I Jungreis, MF Lin, CS Chan, M Kellis 2016). Examination of the alignment can reveal many red flags. If the number of aligned species or their phylogenetic separation is too low, there will be inadequate statistical power to get a meaningful PhyloCSF score. The distribution of species is also important, as alignment to distant species but not closer ones usually indicates non-syntenic alignment of a pseudogene to a paralog. The alignment can be checked for conservation of the ORF itself, namely the start codon, stop codon, and reading frame, without in-frame premature stop codons. (Data in Mackowiak et al. Table S1 indicate that 85 of their ORFs were not conserved in this way in any other species, and another 94 were not conserved beyond gorilla.) The alignment can also be examined to determine if only part of the ORF appears to have evolved as a protein-coding region, indicating that the signature is due to a different splice variant, as well as whether the protein-coding signature appears to continue upstream of the start codon.

In summary, although the Mackowiak et al. smORF predictions proved to be a fruitful ranked list of candidates for discovering novel loci, most of the candidates were not smORFs, limiting their usefulness for discerning smORFs properties.

I Jungreis, MF Lin, CS Chan, M Kellis. 2016. CodAlignView. *CodAlignView: The Codon Alignment Viewer.* https://data.broadinstitute.org/compbio1/cav.php (Accessed April 30, 2016).

Lin MF. 2012. PhyloCSF's "Omega Test." *PhyloCSF's "Omega Test."* http://mlin.github.io/PhyloCSF/OmegaTest.pdf (Accessed June 19, 2021).

Lin MF, Jungreis I, Kellis M. 2011. PhyloCSF: a comparative genomics method to distinguish protein coding and non-coding regions. *Bioinformatics* **27**: i275–82. http://dx.doi.org/10.1093/bioinformatics/btr209.

Mackowiak SD, Zauber H, Bielow C, Thiel D, Kutz K, Calviello L, Mastrobuoni G, Rajewsky N, Kempa S, Selbach M, et al. 2015. Extensive identification and analysis of conserved small ORFs in animals. *Genome Biol* **16**: 179. http://dx.doi.org/10.1186/s13059-015-0742-x.

Mudge JM, Jungreis I, Hunt T, Gonzalez JM, Wright JC, Kay M, Davidson C, Fitzgerald S, Seal R, Tweedie S, et al. 2019. Discovery of high-confidence human protein-coding genes and exons by whole-genome PhyloCSF helps elucidate 118 GWAS loci. *Genome Res* **29**: 2073–2087. http://dx.doi.org/10.1101/gr.246462.118.

Values calculated using ranksum function in MATLAB 2021 (the Wilcoxon rank sum test, a test equivalent to the Mann-Whitney U-test).

| | A | R | N | D | C | Q | E |
|---|---|---|---|---|---|---|---|
| NR-SEPs vs. Negative Control | 1.79E-01 | 4.25E-02 | 2.79E-04 | 1.91E-02 | 4.65E-03 | 5.75E-01 | 7.96E-01 |
| NR-SEPs vs. All Uniprot | 5.90E-05 | 3.63E-02 | 5.61E-04 | 5.91E-16 | 8.00E-02 | 6.86E-03 | 3.23E-13 |
| NR-SEPs vs. Random Control | 7.37E-01 | 1.98E-01 | 5.80E-01 | 6.45E-01 | 1.08E-02 | 5.36E-01 | 1.23E-01 |

| | G | H | I | L | K | M | F |
|---|---|---|---|---|---|---|---|
| NR-SEPs vs. Negative Control | 7.45E-07 | 4.74E-06 | 2.56E-03 | 3.40E-01 | 4.27E-11 | 5.30E-04 | 5.21E-04 |
| NR-SEPs vs. All Uniprot | 7.33E-05 | 4.01E-10 | 8.42E-01 | 7.97E-01 | 2.25E-04 | 1.53E-29 | 9.30E-03 |
| NR-SEPs vs. Random Control | 9.27E-03 | 8.23E-01 | 4.29E-01 | 4.71E-01 | 1.60E-01 | 6.32E-01 | 9.45E-01 |

| | P | S | T | W | Y | V |
|---|---|---|---|---|---|---|
| NR-SEPs vs. Negative Control | 2.44E-11 | 1.89E-08 | 1.27E-01 | 2.35E-03 | 4.03E-07 | 6.74E-01 |
| NR-SEPs vs. All Uniprot | 3.36E-08 | 2.93E-07 | 4.82E-10 | 2.87E-02 | 8.11E-01 | 1.22E-03 |
| NR-SEPs vs. Random Control | 3.12E-01 | 8.84E-01 | 6.92E-01 | 8.64E-01 | 1.06E-02 | 4.42E-01 |

| | Cysteine density | Electrostatic charge density | Net charge density | Length alpha helix | fraction alpha helix | Length of beta strand | Fraction beta strand | Predicted disorder | Low complexity | GRAVY |
|---|---|---|---|---|---|---|---|---|---|---|
| | p value | | | | | | | | | |
| Kruskall-Wallis multiple comparisons test | <0.0001 | <0.0001 | <0.0001 | <0.0001 | <0.0001 | 0.0783 | <0.0001 | <0.0001 | <0.0001 | 0.0001 |
| indicating the dominance of one of the sets | | | | | | | | | | |

subsequent multiple comparisons using the grouped rankings:

| Dunn's multiple comparisons test | Adjusted P Value | | | | | | | | | |
|---|---|---|---|---|---|---|---|---|---|---|
| NR-SEPs vs. Negative Control | <0.0001 | <0.0001 | >0.9999 | <0.0001 | <0.0001 | 0.4721 | <0.0001 | 0.2747 | <0.0001 | >0.9999 |
| NR-SEPs vs. Random Control | <0.0001 | >0.9999 | >0.9999 | <0.0001 | <0.0001 | 0.184 | <0.0001 | 0.0093 | <0.0001 | >0.9999 |
| NR-SEPs vs. All Uniprot or Uniprot-1 | 0.5266 | >0.9999 | <0.0001 | <0.0001 | 0.0245 | <0.0001 | <0.0001 | <0.0001 | 0.1097 | 0.1885 |
| Negative Control vs. Random Control | >0.9999 | <0.0001 | 0.2196 | 0.5265 | 0.0008 | >0.9999 | 0.251 | <0.0001 | 0.0025 | >0.9999 |
| Negative Control vs. All Uniprot or Uniprot-1 | <0.0001 | <0.0001 | <0.0001 | <0.0001 | <0.0001 | 0.0014 | >0.9999 | <0.0001 | <0.0001 | 0.0457 |
| Random Control vs. All Uniprot or Uniprot-1 | <0.0001 | 0.2932 | <0.0001 | <0.0001 | 0.3633 | 0.0039 | 0.0062 | 0.5449 | <0.0001 | 0.014 |

Individual comparisons with Mann-Whitney test

| | P Value | | | | | | | | | |
|---|---|---|---|---|---|---|---|---|---|---|
| NR-SEPs vs. Negative Control | 0.0046 | <0.0001 | 0.0378 | <0.0001 | <0.0001 | 0.0605 | <0.0001 | 0.03008956 | <0.0001 | 0.95526928 |
| NR-SEPs vs. Random Control | 0.0108 | 0.6372 | 0.6803 | <0.0001 | <0.0001 | 0.0264 | <0.0001 | 0.00061068 | <0.0001 | 0.96467932 |
| NR-SEPs vs. All Uniprot or Uniprot-1 | 0.0801 | 0.5494 | <0.0001 | <0.0001 | 0.0008 | <0.0001 | <0.0001 | 1.97E-06 | 0.6027 | 0.03217203 |
| Negative Control vs. Random Control | 0.6262 | <0.0001 | 0.0049 | 0.0722 | <0.0001 | 0.6468 | 0.0268 | 6.27E-08 | <0.0001 | 0.94997883 |
| Negative Control vs. All Uniprot or Uniprot-1 | <0.0001 | <0.0001 | <0.0001 | <0.0001 | <0.0001 | 0.0002 | 0.1177 | 1.39E-10 | <0.0001 | 0.00762857 |
| Random Control vs. All Uniprot or Uniprot-1 | <0.0001 | 0.0484 | <0.0001 | <0.0001 | 0.0521 | 0.0007 | 0.0001 | 0.02707507 | <0.0001 | 0.00225273 |

Individual comparisons with Fisher's Exact test for binary classifications

P values

| | Signal Peptide | Transmembrane Helix | Phase Separation |
|---|---|---|---|
| NR-SEPs vs. Negative Control | 4.11E-05 | 1.21E-14 | 8.17E-05 |
| NR-SEPs vs. Random Control | 3.67E-04 | 1.21E-14 | 0.684001769 |
| NR-SEPs vs. Uniprot-1 | 0.4702 | 0.69161299 | |
| Negative Control vs. Random Control | 1 | 1 | 4.08E-06 |
| Negative Control vs. Uniprot-1 | 9.19E-07 | 3.96E-13 | |
| Random Control vs. Uniprot-1 | 1.08E-05 | 3.96E-13 | |

Table S1: Results from Statistical Tests, as indicated.

| | | | | | | | |
|---|---|---|---|---|---|---|---|
| Analysis Type: | PANTHER Overrepresentation Test (Released 20231017) | | | | | | |
| Annotation Version and Release Date: | PANTHER version 18.0 Released 2023-08-01 | | | | | | |
| Analyzed List: | NR-SEPs (Homo sapiens ) | | | | | | |
| Reference List: | Homo sapiens (all genes in database) | | | | | | |
| Test Type: | FISHER | | | | | | |
| Correction: | FDR | | | | | | |
| PANTHER GO-Slim Cellular Component | Homo sapiens - REFLIST (20592) | NR-SEPs (99) | NR-SEPs (expected) | NR-SEPs (over/under) | NR-SEPs (fold Enrichment) | NR-SEPs (raw P-value) | NR-SEPs (FDR) |
| mitochondrial outer membrane translocase complex (GO:0005742) | 9 | 2 | 0.04 | + | 46.22 | 1.21E-03 | 2.38E-02 |
| mitochondrial proton-transporting ATP synthase complex (GO:0005753) | 14 | 3 | 0.07 | + | 44.57 | 6.88E-05 | 2.82E-03 |
| RNA polymerase II, core complex (GO:0005665) | 10 | 2 | 0.05 | + | 41.6 | 1.45E-03 | 2.74E-02 |
| RNA polymerase I complex (GO:0005736) | 10 | 2 | 0.05 | + | 41.6 | 1.45E-03 | 2.64E-02 |
| proton-transporting ATP synthase complex (GO:0045259) | 17 | 3 | 0.08 | + | 36.71 | 1.14E-04 | 4.01E-03 |
| proton-transporting two-sector ATPase complex (GO:0016469) | 29 | 3 | 0.14 | + | 21.52 | 4.76E-04 | 1.07E-02 |
| respirasome (GO:0070469) | 58 | 5 | 0.28 | + | 17.93 | 1.28E-05 | 5.71E-04 |
| mitochondrial inner membrane (GO:0005743) | 122 | 10 | 0.59 | + | 17.05 | 7.62E-10 | 1.25E-07 |
| organelle inner membrane (GO:0019866) | 129 | 10 | 0.62 | + | 16.12 | 1.27E-09 | 1.56E-07 |
| mitochondrial respirasome (GO:0005746) | 52 | 4 | 0.25 | + | 16 | 1.50E-04 | 4.60E-03 |
| respiratory chain complex (GO:0098803) | 53 | 4 | 0.25 | + | 15.7 | 1.60E-04 | 4.64E-03 |
| mitochondrial membrane (GO:0031966) | 177 | 12 | 0.85 | + | 14.1 | 1.10E-10 | 5.41E-08 |
| inner mitochondrial membrane protein complex (GO:0098800) | 89 | 6 | 0.43 | + | 14.02 | 6.35E-06 | 3.12E-04 |
| mitochondrial envelope (GO:0005740) | 193 | 12 | 0.93 | + | 12.93 | 2.82E-10 | 6.93E-08 |
| mitochondrial protein-containing complex (GO:0098798) | 134 | 8 | 0.64 | + | 12.42 | 4.08E-07 | 2.87E-05 |
| cytosolic ribosome (GO:0022626) | 85 | 4 | 0.41 | + | 9.79 | 8.82E-04 | 1.81E-02 |
| envelope (GO:0031975) | 291 | 12 | 1.4 | + | 8.58 | 2.32E-08 | 2.29E-06 |
| organelle envelope (GO:0031967) | 291 | 12 | 1.4 | + | 8.58 | 2.32E-08 | 1.91E-06 |
| mitochondrion (GO:0005739) | 602 | 14 | 2.89 | + | 4.84 | 1.37E-06 | 8.44E-05 |
| organelle membrane (GO:0031090) | 733 | 15 | 3.52 | + | 4.26 | 2.62E-06 | 1.43E-04 |
| membrane protein complex (GO:0098796) | 653 | 11 | 3.14 | + | 3.5 | 3.33E-04 | 8.20E-03 |
| Unclassified (UNCLASSIFIED) | 8704 | 60 | 41.85 | + | 1.43 | 3.15E-04 | 8.60E-03 |
| cellular anatomical entity (GO:0110165) | 11637 | 39 | 55.95 | - | 0.7 | 7.36E-04 | 1.58E-02 |
| cellular_component (GO:0005575) | 11888 | 39 | 57.15 | - | 0.68 | 3.15E-04 | 8.15E-03 |
| nucleus (GO:0005634) | 3516 | 4 | 16.9 | - | 0.24 | 1.34E-04 | 4.40E-03 |
| plasma membrane (GO:0005886) | 2192 | 1 | 10.54 | - | 0.09 | 4.11E-04 | 9.63E-03 |
| cell periphery (GO:0071944) | 2518 | 1 | 12.11 | - | 0.08 | 7.37E-05 | 2.79E-03 |

Table S2: PANTHER cellular component statistical overrepresentation of the 100 NR-SEPs annotated in the database, as compared to the human proteome. The columns are: 1) The PANTHER GO-Slim Biological Process ontology terms, 2) The number of proteins in the human proteome reference with that ontology term, 3) the number of proteins in the NR-SEP list annotated with that term (note that only 100 of the proteins have ontology terms associated with their UniProt IDs), 4) the fraction of proteins with that annotation term in the reference human proteome, 5) the sign of enrichment of SEPs vs the human proteome, 6) the fold-enrichment of SEPs with that annotation term relative to the that of the human proteome, 7) the p-value and 8) the false discovery rate.

| | | | | | | | |
|---|---|---|---|---|---|---|---|
| Analysis Type: | PANTHER Overrepresentation Test (Released 20231017) | | | | | | |
| Annotation Version and Release Date: | PANTHER version 18.0 Released 2023-08-01 | | | | | | |
| Analyzed List: | NR-SEPs (Homo sapiens ) | | | | | | |
| Reference List: | Homo sapiens (all genes in database) | | | | | | |
| Test Type: | FISHER | | | | | | |
| Correction: | FDR | | | | | | |
| PANTHER GO-Slim Biological Process | Homo sapiens - REFLIST (20592) | NR-SEPs (99) | NR-SEPs (expected) | NR-SEPs (over/under) | NR-SEPs (fold Enrichment) | NR-SEPs (raw P-value) | NR-SEPs (FDR) |
| oxidative phosphorylation (GO:0006119) | 27 | 3 | 0.13 | + | 23.11 | 3.93E-04 | 3.60E-02 |
| defense response to bacterium (GO:0042742) | 61 | 4 | 0.29 | + | 13.64 | 2.67E-04 | 2.94E-02 |
| Unclassified (UNCLASSIFIED) | 8302 | 73 | 39.91 | + | 1.83 | 2.07E-11 | 2.27E-08 |
| cellular process (GO:0009987) | 7929 | 19 | 38.12 | - | 0.5 | 4.51E-05 | 9.04E-03 |
| biological_process (GO:0008150) | 12290 | 26 | 59.09 | - | 0.44 | 2.07E-11 | 4.55E-08 |
| biological regulation (GO:0065007) | 6092 | 11 | 29.29 | - | 0.38 | 1.98E-05 | 7.27E-03 |
| regulation of biological process (GO:0050789) | 5788 | 10 | 27.83 | - | 0.36 | 2.41E-05 | 7.57E-03 |
| metabolic process (GO:0008152) | 3952 | 6 | 19 | - | 0.32 | 2.96E-04 | 3.10E-02 |
| regulation of cellular process (GO:0050794) | 5312 | 7 | 25.54 | - | 0.27 | 2.68E-06 | 1.97E-03 |
| primary metabolic process (GO:0044238) | 3406 | 4 | 16.38 | - | 0.24 | 2.02E-04 | 2.35E-02 |
| organic substance metabolic process (GO:0071704) | 3744 | 4 | 18 | - | 0.22 | 6.01E-05 | 1.10E-02 |
| regulation of nitrogen compound metabolic process (GO:0051171) | 2575 | 2 | 12.38 | - | 0.16 | 3.56E-04 | 3.57E-02 |
| regulation of primary metabolic process (GO:0080090) | 2602 | 2 | 12.51 | - | 0.16 | 3.58E-04 | 3.43E-02 |
| regulation of macromolecule metabolic process (GO:0060255) | 2787 | 2 | 13.4 | - | 0.15 | 1.58E-04 | 1.94E-02 |
| regulation of metabolic process (GO:0019222) | 2939 | 2 | 14.13 | - | 0.14 | 6.71E-05 | 9.86E-03 |
| regulation of nucleobase-containing compound metabolic process (GO:0019219) | 2176 | 0 | 10.46 | - | < 0.01 | 2.41E-05 | 6.64E-03 |
| regulation of macromolecule biosynthetic process (GO:0010556) | 2088 | 0 | 10.04 | - | < 0.01 | 6.26E-05 | 1.06E-02 |
| regulation of transcription by RNA polymerase II (GO:0006357) | 1649 | 0 | 7.93 | - | < 0.01 | 5.34E-04 | 4.71E-02 |
| regulation of DNA-templated transcription (GO:0006355) | 1958 | 0 | 9.41 | - | < 0.01 | 9.14E-05 | 1.26E-02 |
| regulation of gene expression (GO:0010468) | 2409 | 0 | 11.58 | - | < 0.01 | 9.95E-06 | 4.38E-03 |
| regulation of RNA metabolic process (GO:0051252) | 2111 | 0 | 10.15 | - | < 0.01 | 3.74E-05 | 8.24E-03 |
| regulation of biosynthetic process (GO:0009889) | 2115 | 0 | 10.17 | - | < 0.01 | 3.74E-05 | 9.16E-03 |
| regulation of cellular biosynthetic process (GO:0031326) | 2091 | 0 | 10.05 | - | < 0.01 | 6.31E-05 | 9.92E-03 |
| regulation of cellular metabolic process (GO:0031323) | 2536 | 0 | 12.19 | - | < 0.01 | 3.80E-06 | 2.09E-03 |
| regulation of RNA biosynthetic process (GO:2001141) | 1965 | 0 | 9.45 | - | < 0.01 | 9.18E-05 | 1.19E-02 |

Table S3: PANTHER biological processes statistical overrepresentation of the 99 NR-SEPs annotated in the PANTHER database as compared to the human proteome. The columns are: 1) The PANTHER GO-Slim Biological Process ontology terms, 2) The number of proteins in the human proteome reference with that ontology term, 3) the number of proteins in the NR-SEP list annotated with that term (note that only 99 of the proteins have ontology terms associated with their UniProt IDs), 4) the fraction of proteins with that annotation term in the reference human proteome, 5) the sign of enrichment of NR-SEPs vs the human proteome, 6) the fold-enrichment of NR-SEPs with that annotation term relative to the that of the human proteome, 7) the p-value and 8) the false discovery rate.

### Supplemental Figures

A

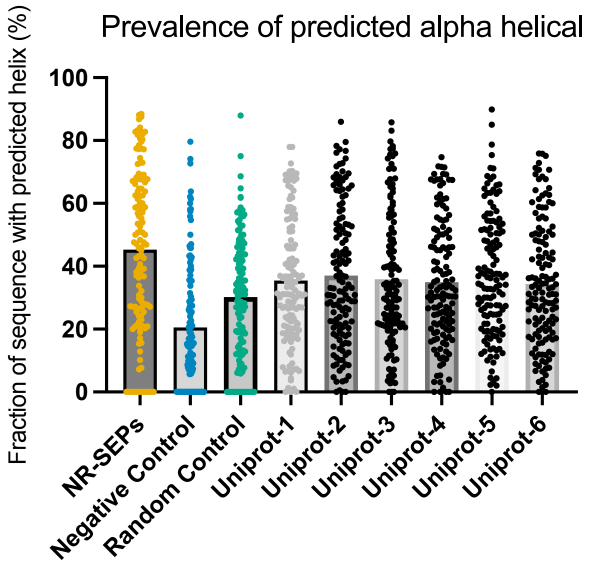


| | NR-SEPs | Negative Control | Random Control | Uniprot-1 |
|---|---|---|---|---|
| **Negative Control** | <0.0001 | | | |
| **Random Control** | <0.0001 | 0.0021 | | |
| **Uniprot-1** | 0.0702 | <0.0001 | >0.9999 | |
| **Uniprot-2** | 0.252 | <0.0001 | 0.7475 | >0.9999 |
| **Uniprot-3** | 0.0588 | <0.0001 | >0.9999 | >0.9999 |
| **Uniprot-4** | 0.0362 | <0.0001 | >0.9999 | >0.9999 |
| **Uniprot-5** | 0.4028 | <0.0001 | 0.4955 | >0.9999 |
| **Uniprot-6** | 0.0149 | <0.0001 | >0.9999 | >0.9999 |

B

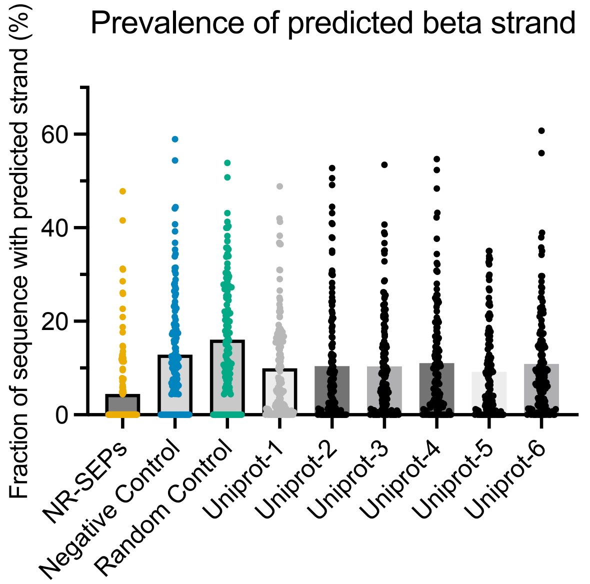


| | NR-SEPs | Negative Control | Random Control | Uniprot-1 |
|---|---|---|---|---|
| **Negative Control** | <0.0001 | | | |
| **Random Control** | <0.0001 | >0.9999 | | |
| **Uniprot-1** | <0.0001 | >0.9999 | 0.0122 | |
| **Uniprot-2** | <0.0001 | >0.9999 | 0.0012 | >0.9999 |
| **Uniprot-3** | <0.0001 | >0.9999 | 0.0287 | >0.9999 |
| **Uniprot-4** | <0.0001 | >0.9999 | 0.0798 | >0.9999 |
| **Uniprot-5** | <0.0001 | >0.9999 | 0.0014 | >0.9999 |
| **Uniprot-6** | <0.0001 | >0.9999 | 0.2378 | >0.9999 |

Figure S1. We used 140 randomly chosen proteins from the Uniprot UP000005640 reference proteome as a comparison to typical well-studied proteins. To investigate whether our data sets were large enough that a single sampling of the reference proteome was sufficient, we redid the analysis of the prevalence of (A) alpha helicies, (B) beta strands for 6 total independent of the reference proteome. Listed p-values are from the Dunn's multiple comparisons test. While there was some variability in the precise p-values, the trends for all analysis were similar independent of reference set. The p-values for all pairwise comparisons between Uniprot sampled sets were all >0.9999.

A

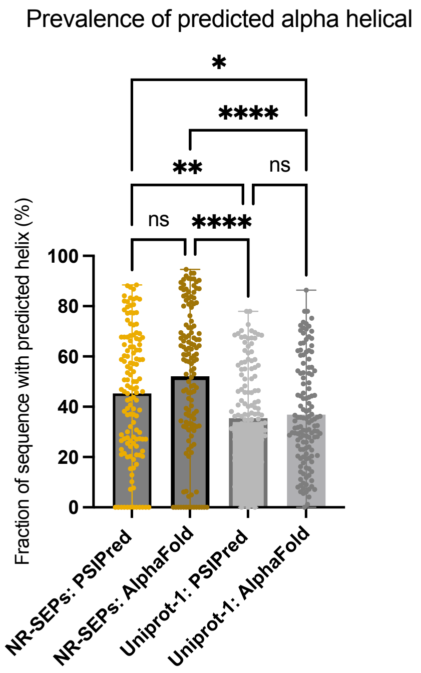


| | NR-SEPs: PSIPred | NR-SEPs: AlphaFold | Uniprot-1: PSIPred |
|---|---|---|---|
| **NR-SEPs: AlphaFold** | 0.1784 | | |
| **Uniprot-1: PSIPred** | 0.0070 | <0.0001 | |
| **Uniprot-1: AlphaFold** | 0.0342 | <0.0001 | >0.9999 |

B

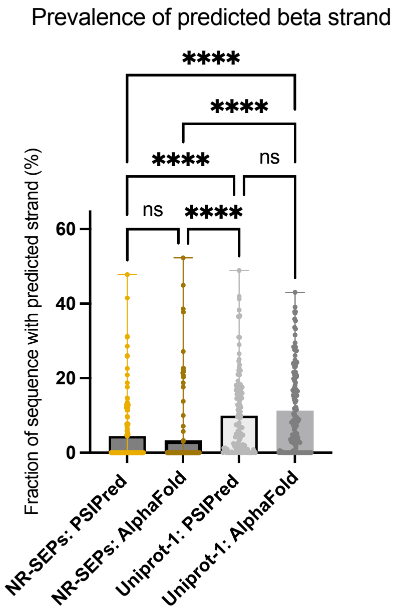


| | NR-SEPs: PSIPred | NR-SEPs: AlphaFold | Uniprot-1: PSIPred |
|---|---|---|---|
| **NR-SEPs: AlphaFold** | 0.2824 | | |
| **Uniprot-1: PSIPred** | <0.0001 | <0.0001 | |
| **Uniprot-1: AlphaFold** | <0.0001 | <0.0001 | >0.9999 |

Figure S2. Secondary structural determination of the prevalence of (A) alpha helicies and (B) beta strands using DSSP to identify secondary structure from Alpha-Fold predicted structures yields very similar results to those found using PSIPred when comparing the NR-SEP and UniProt-1 sets. Beta strands are strongly underrepresented in the NR-SEP dataset. Listed p-values are from the Dunn's multiple comparisons test.

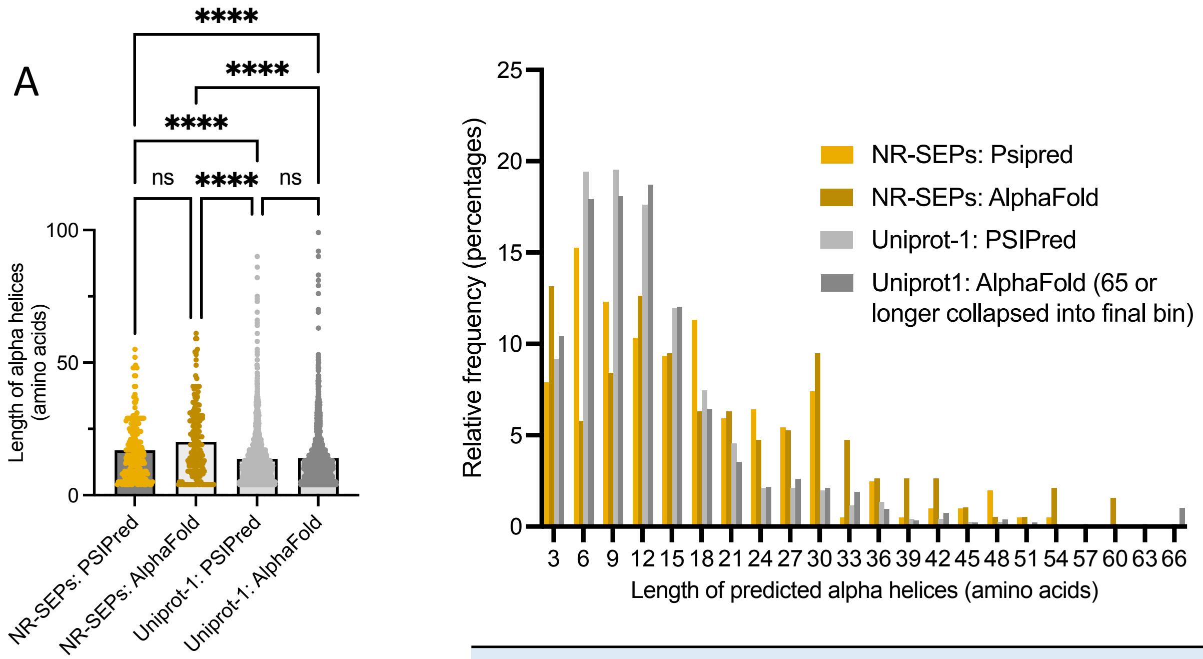


| | NR-SEPs: PSIPred | NR-SEPs : AlphaFold | Uniprot-1: PSIPred |
|---|---|---|---|
| **NR-SEPs: AlphaFold** | 0.4250 | | |
| **Uniprot-1: PSIPred** | <0.0001 | <0.0001 | |
| **Uniprot-1: AlphaFold** | <0.0001 | <0.0001 | >0.9999 |

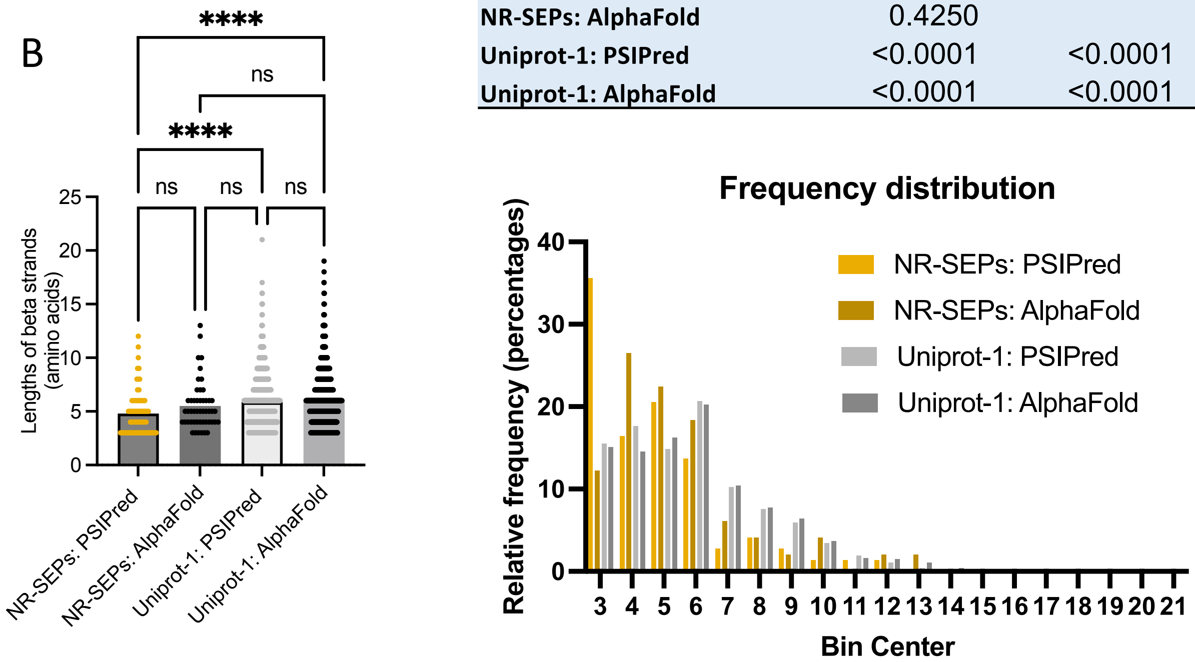


| | NR-SEPs: PSIPred | NR-SEPs: AlphaFold | Uniprot-1: PSIPred |
|---|---|---|---|
| **NR-SEPs: AlphaFold** | 0.3848 | | |
| **Uniprot-1: PSIPred** | <0.0001 | >0.9999 | |
| **Uniprot-1: AlphaFold** | <0.0001 | 0.5347 | 0.3862 |

Figure S3. Secondary structural determination using DSSP to identify secondary structure from Alpha-Fold predicted structures yields very similar results to PSIPred when comparing the length of (A) alpha helicies or (B) beta strands in the NR-SEP and UniProt-1 sets. The NR-SEP set has a higher prevalence of longer alpha helices. AlphaFold predicts slightly more short beta strands as compared to PSIPred which finds more NR-SEPs that lack any bet strands than UniProt-1.

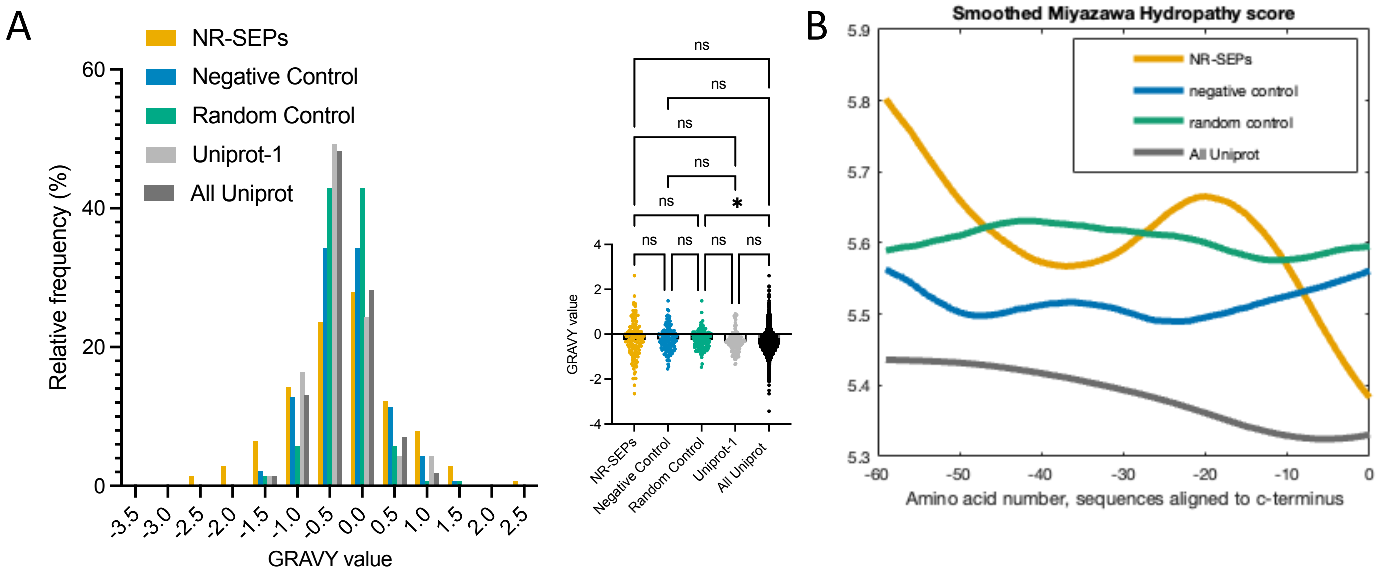


Figure S4. Calculated biophysical characteristics of smORFs. (A) Grand average of hydropathy (GRAVY) was calculated for each sequence. the standard deviation is almost twice as large for the NR-SEPs as for UniProt proteins (0.83 vs 0.43). (B) Hydrophobicity was calculated using the Miyazawa scale of sequences aligned from the C-terminus and smoothed, showing a decrease in hydrophobicity at the C-termini for the UniProt and NR-SEP sets, but not for the controls.

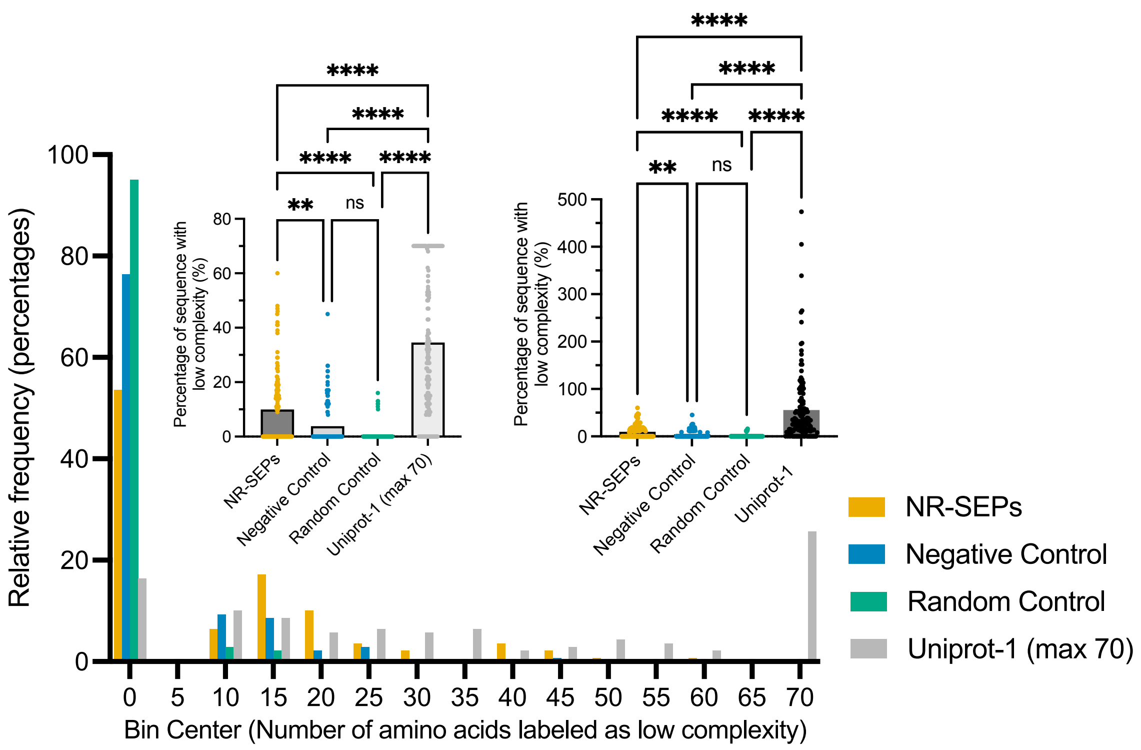


Figure S5. Raw number of amino acids labeled as low complexity. For Uniprot-1 proteins, all proteins with 70 or more amino acids labeled as low complexity were binned into the largest bin.

Figure S6 (continued on next two pages). Comparing the subset of NR-SEPs with experimental evidence of translation in Uniprot to the subset of NR-SEPs that did not have protein function annotated in Uniprot at the time of our query (March 2022). A comparison of the (A) cysteine density, (B) electrostatic charge density, (C) net charge density, (D) predicted low complexity, (E) predicted disorder, (F), predicted beta strand content, (G) predicted alpha helical content, (H) predicted GRAVY scores demonstrates a high level of statistical similarity between the two data subsets. Only the GRAVY scores were normally distributed and so compared using unpaired t-test with Welch's correction. All other datasets were compared using both the Kolmogorov-Smirnov or Mann-Whitney tests. P-values for comparisons were greater than 0.05 by all tests.

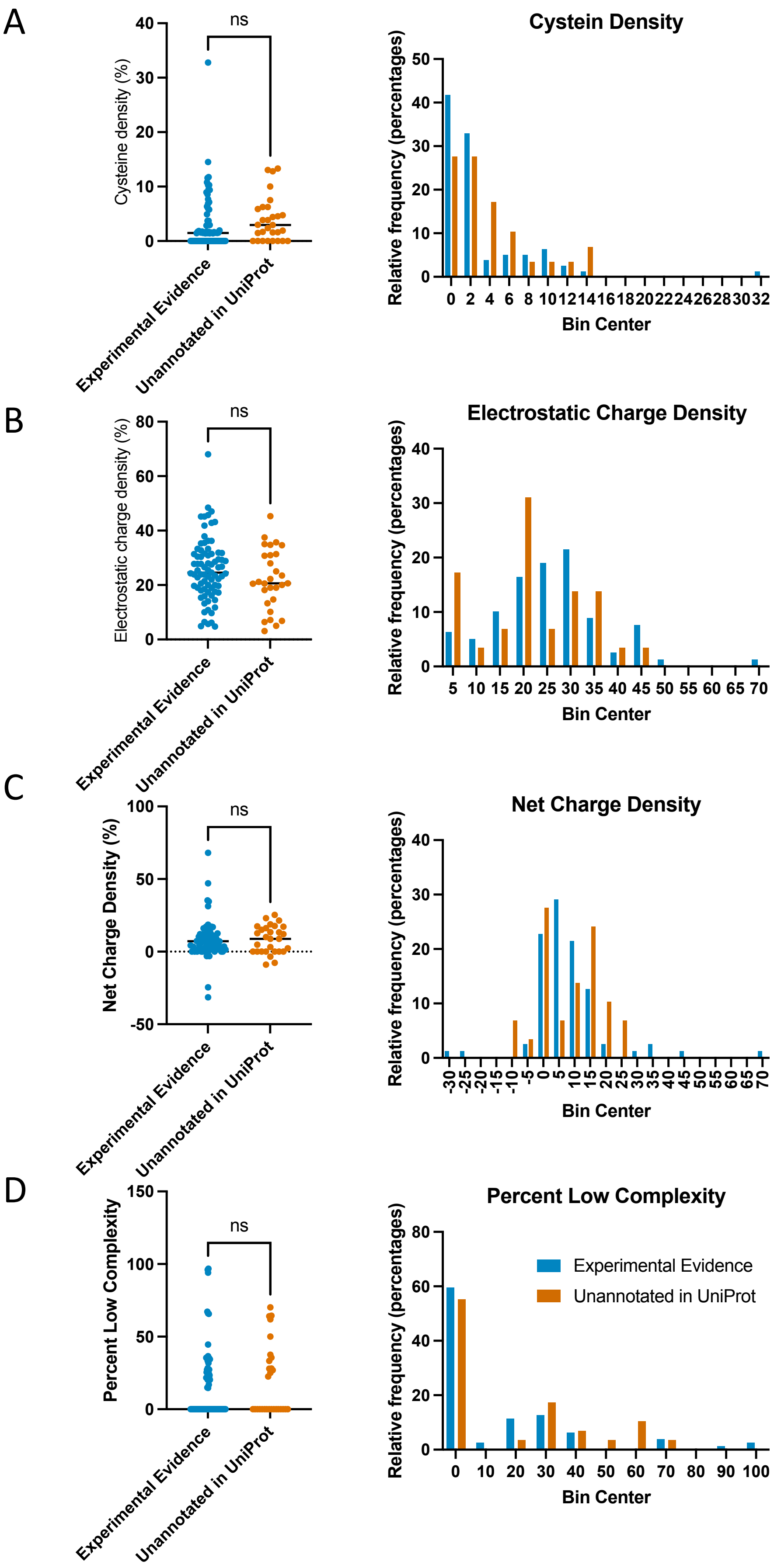

A
ns
Cysteine density (%)
Experimental Evidence
Unannotated in UniProt
Cystein Density
Relative frequency (percentages)
Bin Center
B
ns
Electrostatic charge density (%)
Experimental Evidence
Unannotated in UniProt
Electrostatic Charge Density
Relative frequency (percentages)
Bin Center
C
ns
Net Charge Density (%)
Experimental Evidence
Unannotated in UniProt
Net Charge Density
Relative frequency (percentages)
Bin Center
D
ns
Percent Low Complexity
Experimental Evidence
Unannotated in UniProt
Percent Low Complexity
Experimental Evidence
Unannotated in UniProt
Relative frequency (percentages)
Bin Center

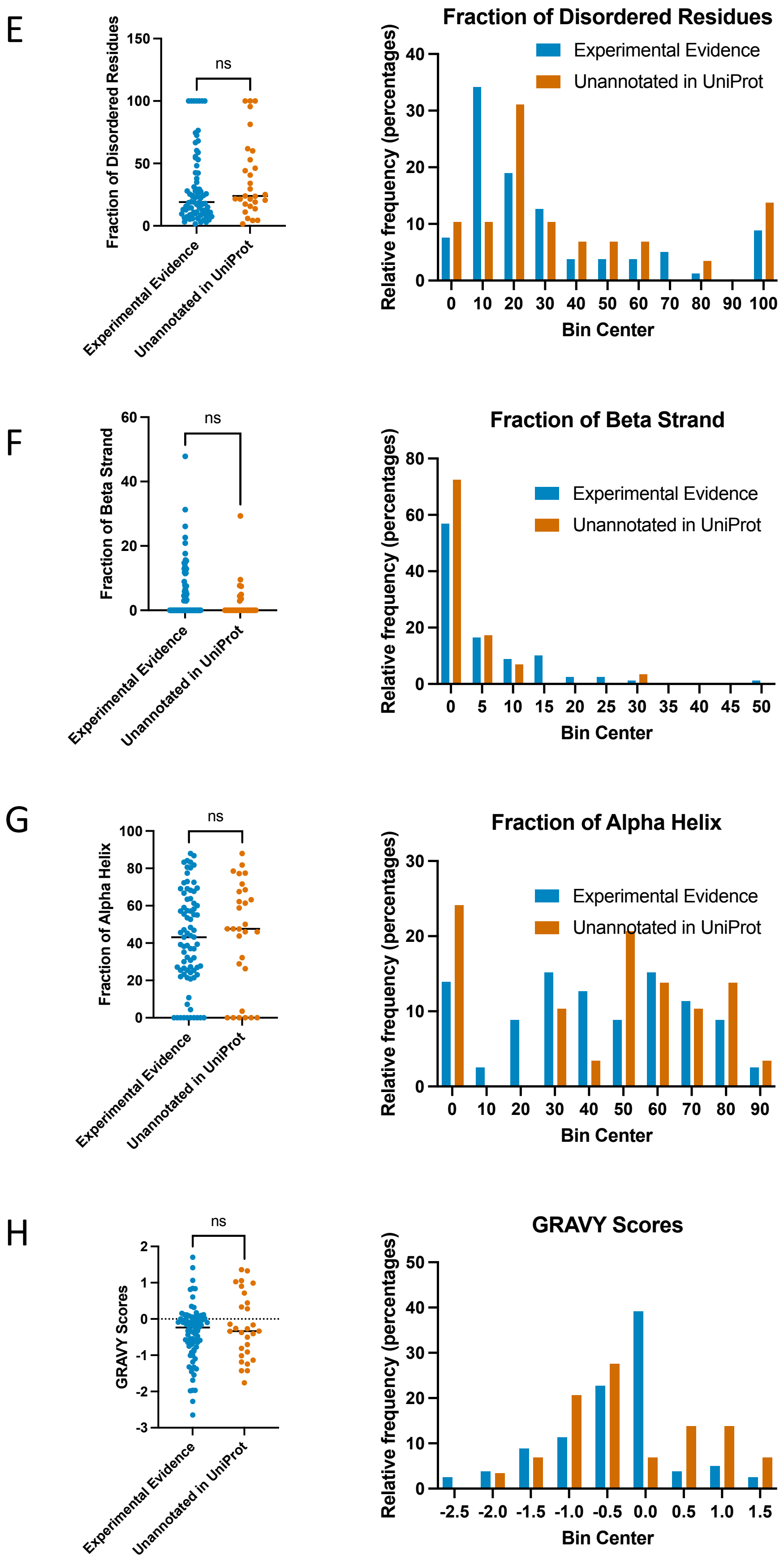
E
Fraction of Disordered Residues
ns
Experimental Evidence
Unannotated in UniProt
Fraction of Disordered Residues
Relative frequency (percentages)
Bin Center
F
Fraction of Beta Strand
ns
Experimental Evidence
Unannotated in UniProt
Fraction of Beta Strand
Relative frequency (percentages)
Bin Center
G
Fraction of Alpha Helix
ns
Experimental Evidence
Unannotated in UniProt
Fraction of Alpha Helix
Relative frequency (percentages)
Bin Center
H
GRAVY Scores
ns
Experimental Evidence
Unannotated in UniProt
GRAVY Scores
Relative frequency (percentages)
Bin Center
